# Nuclear size is genetically controlled and influences cell fate

**DOI:** 10.64898/2026.08.26.747275

**Authors:** Hisashi Moriizumi, Ruyi Shi, Daniel Schraivogel, Adam James Reid, Lars M. Steinmetz, Jan M. Skotheim, Evgeny Zatulovskiy

## Abstract

The proportional scaling between nuclear and cell size was first described more than 150 years ago and is among the most conserved features of cellular organization. Yet the mechanisms that establish this scaling and its physiological significance have remained unresolved. Here, we address both questions by combining image-enabled cell sorting with genome-wide CRISPR screening, transcriptomics and functional analyses. We identify more than 180 regulators of the nuclear-to-cell ratio and show that distinct classes of genes independently control nuclear and cell size. RNA metabolism predominantly regulates nuclear size, whereas protein synthesis and degradation primarily regulate cell size. This supports a model in which differences in macromolecular partitioning between the nucleus and cytoplasm contribute to osmotic regulation of nuclear size together with mechanical constraints imposed by chromatin, the cytoskeleton, and the nuclear envelope. Changes in nuclear size cause widespread transcriptional remodeling that is independent of changes in cell size. Cells with smaller nuclei exhibit reduced PRC2-dependent H3K27 trimethylation, activation of developmental gene-expression programmes and repression of cell-cycle genes. Consistent with these changes, mouse embryonic stem cells with smaller nuclei show an increased propensity to exit pluripotency and initiate differentiation in response to retinoic acid. Together, our findings provide a mechanistic framework for nuclear size scaling and establish nuclear size as a physical regulator of gene expression and cell-state transitions, linking cellular architecture to cell fate.

## Introduction

Cell size is a fundamental feature of cell geometry that sets the scale for many intracellular structures including the mitochondria, vacuole, spindle and nucleus^1–6^. The best characterized example of this phenomenon is that the nuclear volume always represents a fixed fraction of the cell volume for a given cell type, which was first reported in an 1870’s study comparing cellular and nuclear sizes of red blood cells across a variety of vertebrate species^7,8^. The constancy of the nuclear-to-cellular volume ratio (N/C ratio) was so striking that researchers in the early 1900’s gave it the special name of karyoplasmic ratio^9^. Extensive cytological studies over the past century revealed that the N/C ratio is tightly controlled across diverse organisms and cell types, with nuclear size scaling in direct proportion to cell size^1,10^. For example, in fission yeast it was shown that the N/C ratio remains constant over a 35-fold range of cell sizes and is maintained independently of cell shape, highlighting the remarkable robustness of N/C scaling mechanisms^1,10^. The evolutionary conservation and robustness of this proportional scaling relationship suggest that nuclear size is fundamentally important for cell physiology and tightly coupled to cell volume. However, how and why nuclear size is controlled still remains a fundamental unresolved question in cell biology (**Fig. 1a**).

**Fig. 1.**
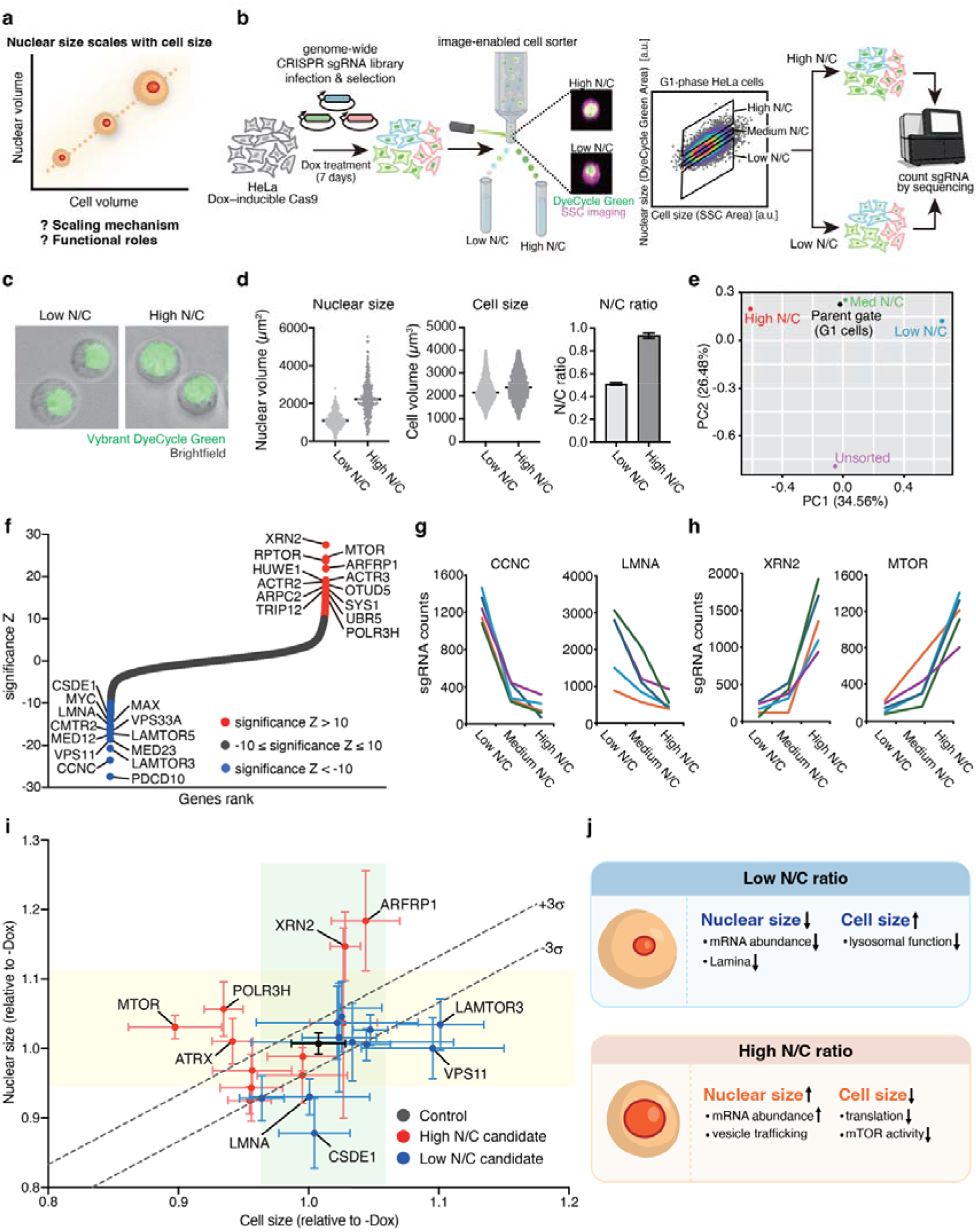
Genome-wide CRISPR screen identifies genes affecting the N/C ratio in human cells. **a,** A schematic illustrating the proportional scaling between nuclear volume and cell volume. **b**, Schematic overview of the imaging flow cytometry-based genome-wide CRISPR knockout screen. Tet::Cas9 HeLa cells were transduced with a pooled sgRNA library and treated with doxycycline for 7 days, after which cells were harvested, stained with Vybrant DyeCycle Green, and sorted using an image-enabled cell sorter into low, medium, and high N/C ratio populations based on nuclear and cell area measurements. Representative images show nuclear fluorescence (green) and cell morphology (magenta), visualized by Vybrant DyeCycle Green staining and side scatter (SSC) imaging, respectively. **c,** Representative widefield microscopy images of sorted low N/C and high N/C cells, showing nuclear fluorescence (Vybrant DyeCycle Green) overlaid with brightfield images. **d,** Quantification of nuclear volume (left), cell volume (middle), and N/C ratio (right) in cells sorted into low- and high-N/C ratio populations. Nuclear images were acquired using Vybrant DyeCycle Green DNA staining. Each dot represents an individual cell. Bars indicate mean ± SEM. **e**, Principal component analysis (PCA) of sgRNA abundance profiles in unsorted cells and N/C-sorted G1 populations of HeLa cells. The low-, medium-, and high-N/C-ratio populations appear ordered along PC1 according to their N/C ratios. Each point represents the sgRNA abundance profile of the indicated population, averaged across five biological replicates (i.e., five independent genome-wide CRISPR libraries). **f**, Genes whose sgRNA enrichment significantly increased or decreased with N/C ratio, as determined using the MAUDE algorithm^35^. Each point represents a gene plotted according to its significance Z score; selected high-confidence hits (Z < -10 or Z > 10) are indicated. **g**, **h**, Normalised sgRNA read counts for representative genes enriched in low (g) and high (h) N/C ratio populations. Data are shown for five independent biological replicates, each using a different genome-wide sgRNA sublibrary. **i**, Validation of high-ranked N/C-affecting genes identified in the CRISPR screen. Changes in nuclear size (y-axis) and cell size (x-axis) in Dox-induced knockout cell lines were calculated relative to the corresponding uninduced cells. Red and blue symbols indicate candidate genes originally identified in the screen as enriched in high- and low-N/C ratio populations, respectively, and black symbols indicate the non-targeting control sgRNA. Nuclear size was measured with microscopy, and cell volume was measured on a Coulter counter. Each point represents the mean of n = 3–5 independent experiments, and error bars indicate standard deviation (SD). Dashed lines indicate thresholds corresponding to a constant N/C ratio ± 3σ (σ = SD of the non-targeting control sgRNA). Genes primarily affecting nuclear size are highlighted with green shading, while genes primarily affecting cell size are highlighted with yellow. **j**, A schematic summary of genetic factors affecting the N/C ratio by changing either the nuclear size or cell size, according to the CRISPR screen and validation experiments.

Nuclear size control is likely to be important for gene regulation and cell-state transitions because the N/C ratio is known to undergo well-characterized changes during development and differentiation. For example, pluripotent stem cells typically exhibit a high N/C ratio and large nuclear size^11,12^. Nuclear size then decreases during differentiation, as was demonstrated in several stem-cell models, including neural and adipogenic differentiation protocols^13–15^. However, whether the reduction in nuclear size that accompanies differentiation is simply a morphological consequence of cell-state change or actively contributes to the transition from pluripotency to differentiation remains unknown. In addition, the N/C ratio is frequently altered in cancer, where enlarged nuclei relative to cell size are a hallmark of malignant cells and are associated with tumour progression^16^. Despite these well-documented observations, it remains unclear whether changes in the N/C ratio actively influence gene expression programmes and cellular physiology, or instead simply reflect underlying differences in cell type, developmental state, or disease status.

Nuclear morphology can influence cellular physiology by altering nuclear architecture, chromatin organization and the biophysical properties of the nucleoplasm, thereby modulating gene expression. For example, a study in fission yeast has shown that experimentally altering nuclear size can affect the rheological properties of the nucleoplasm and tune heterochromatin condensation^17^. Perturbations of nuclear elongation in early *Drosophila* embryos have been associated with altered heterochromatin organization and changes in gene expression during early embryonic development^18^. Increased nuclear size in human pluripotent stem cells has also been found to promote the formation of nuclear condensates containing the lncRNA *NEAT1*, which regulate differentiation^19^. In epithelial cells, constraining the nucleus leads to downregulation of the euchromatic mark H3K9ac and upregulation of the heterochromatic mark H3K27me3^20^. These findings raise the possibility that nuclear geometry, including nuclear size, is not just a passive scaling property, but a regulatory parameter that actively shapes global gene expression programmes and cell-state transitions.

How the N/C ratio is maintained also remains unknown. Previous studies using gene knockouts and biochemical reconstitution systems identified several classes of genes and cellular processes that influence the N/C ratio, including genes regulating chromatin organization, nuclear lamina structure, nucleocytoplasmic transport, and membrane dynamics, such as lipid supply^6,21–24^. However, it remains unclear whether these mechanisms are responsible for coupling nuclear size to cell size in wild-type cells under physiological conditions. Studies in rapidly dividing *Xenopus* embryos have shown that nucleocytoplasmic transport can influence nuclear growth during short embryonic cell cycles. However, these embryos represent a highly dynamic, non-steady-state system, and it remains unclear whether similar mechanisms maintain nuclear size in steady-state mammalian cells. Systematic studies in fission yeast have dissected N/C ratio regulation by combining mutant screens with quantitative measurements of nuclear and cell size^25,26^. These screens identified multiple cellular processes involved in nuclear size control, including RNA processing and nucleocytoplasmic mRNA transport, lipid metabolism linked to nuclear membrane growth, and LINC-complex-mediated coupling between the nucleus and cytoskeleton. However, the magnitude of N/C ratio alterations observed in these screens was modest (on the order of 10%), and the list of identified mutants did not immediately point to a particular molecular mechanism regulating the N/C ratio. In addition to these genetic approaches, another line of research has focused on the biophysical models of N/C ratio regulation in fission yeast, building on a colloid osmotic framework for intracellular crowding^27,28^. This work suggested that differential partitioning of macromolecules between the nucleoplasm and cytoplasm generates osmotic forces on the nuclear envelope that contribute to nuclear size control^29^. While this model well described the nuclear size regulation in fission yeast, which has little nuclear tension, it is not clear to what extent it is applicable to mammalian cells, whose nuclei are mechanically supported by lamins^30^, which also contribute to tethering chromatin to the nuclear periphery and cellular cytoskeleton^31^. Chromatin itself can also contribute to nuclear stiffness and morphology^32^. By contrast, fission yeast lacks lamins and therefore lack this lamin-based layer of nuclear mechanical regulation. Fission yeast also possesses a cell wall, and their cellular dimensions are mechanically constrained by the balance between turgor pressure and cell-wall resistance^29^. It therefore remains unclear whether osmotic effects arising from macromolecular partitioning are sufficient to explain regulation of the N/C ratio in mammalian cells or whether they act in concert with structural and mechanical determinants of nuclear size. In mammalian cells, systematic analyses of N/C ratio regulation have been limited, largely due to the lack of scalable methods for simultaneously quantifying nuclear and cell size at single-cell resolution across many genetic perturbations.

In this work, we address the regulation of nuclear size in human cells by taking advantage of cutting-edge image-enabled cell sorting technology that enables simultaneous quantification of nuclear and cell size at single-cell resolution and at scales compatible with genetic screening^33^. We combine this approach with RNA sequencing and genome-wide CRISPR screening to characterize N/C-dependent changes in gene expression in human cells and identify the molecular pathways controlling the N/C ratio. We demonstrate that changes in nuclear size affect the expression of developmental genes and genes involved in epigenetic regulation, including components of Polycomb complexes. Moreover, we show that among mouse embryonic stem cells (mESCs) of similar cell size, those with smaller nuclei more readily exit pluripotency and initiate neural differentiation, whereas those with larger nuclei are more likely to retain pluripotency. Given that nuclear size typically decreases during mammalian stem cell differentiation, our results suggest that nuclear-size reduction may not simply be a morphological consequence of differentiation, but it may be actively involved in regulating the differentiation-associated transcriptional programmes and modulating propensity of stem cells to exit pluripotency.

Our genome-wide CRISPR screen reveals that the N/C ratio in human cells is modulated by genes regulating RNA and protein metabolism, mitochondrial activity, the nuclear lamina, the actin cytoskeleton and chromatin organization. These findings support a mechano-osmotic model in which RNA- and protein-dependent osmotic pressures integrate with structural forces to tune nuclear and cell size, linking cellular geometry to gene expression and cell state. Taken together, our work makes substantial progress on a fundamental problem in cell biology that has remained unresolved for more than a century, demonstrating that nuclear size is a key regulator of gene expression and cell-state transitions, and that the N/C ratio emerges from the collective action of distinct molecular pathways independently controlling nuclear and cell size.

## Results

### Genome-wide CRISPR screen identifies genes affecting the N/C ratio in human cells

To identify the molecular mechanisms regulating the N/C ratio in mammalian cells, we first sought to systematically identify genes whose expression affects the N/C ratio in human cells. To achieve this, we combined the power of high-speed image-enabled cell sorting (ICS) technology^33^ with genome-wide CRISPR/Cas9-mediated knockout screen. ICS enabled us to visualize both the cells and their nuclei in a high-throughput flow cytometry platform so that we could isolate cell populations based on their N/C ratio. For this screen, we transduced HeLa cells expressing doxycycline (Dox)–inducible Cas9 (HeLa Tet::Cas9 cell line cTT20^34^) with genome-wide lentiviral single-guide RNA (sgRNA) libraries targeting 18,408 protein-coding genes^33^. This screen was performed using five independent genome-wide sgRNA libraries. We induced CRISPR/Cas9-mediated knockouts by adding Dox to the cells and then performed the N/C-based sorting 7 days after knockout induction. For the sorting, we labelled the cells with a DNA-staining green fluorescent dye Vybrant DyeCycle Green and used ICS to isolate G1-phase cells with high, medium, and low N/C ratios. Each population contained about 2.2 million cells, a scale compatible with genome-wide screening (**Fig. 1b** and **Fig. S1a, b**). After sorting, we quantified nuclear and cell volumes using widefield fluorescence microscopy and a Coulter counter, respectively, confirming that average nuclear size differed, while cell size remained similar across the sorted cell population, resulting in an approximately 1.8-fold range in N/C ratios (**Fig. 1c, d** and **Fig. S1c**).

After isolating cells with high, medium, and low N/C ratios, we used amplicon sequencing to identify sgRNAs enriched in those cell populations (**Supplementary Data 1**). Principal component analysis (PCA) of sgRNA abundance profiles showed that the high-, medium-, and low-N/C-ratio populations were ordered along PC1 according to their N/C ratios (**Fig. 1e**). This alignment indicates that the N/C ratio was a key determinant of sgRNA composition in the sorted cell populations. To identify specific genes whose sgRNAs were enriched in low- or high-N/C ratio populations we used MAUDE statistical analysis^35^ to calculate Z scores based on sgRNA read counts. Positive Z scores correspond to sgRNA enrichment in the high N/C-ratio population, whereas negative Z scores correspond to enrichment in the low N/C-ratio population. Using thresholds of Z > 10 and Z < −10, we identified 137 genes whose sgRNAs were enriched in the high N/C population of cells, and 52 genes whose sgRNAs were enriched in the low N/C population of cells (**Fig. 1f** and **Supplementary Data 2**). The results for these genes were consistent across the five different genome-wide libraries used in the experiment (**Fig. 1g, h**). Among the identified hits, we recovered known regulators of nuclear and cell size, such as *LMNA* and *MTOR*, whose loss has been reported to reduce nuclear size and cell size, respectively^22,36^ (**Fig. 1 g, h**), as well as previously uncharacterized and unexpected candidates.

To validate the hits identified in the genome-wide screen, we selected 22 highly ranked candidate genes with distinct biological functions, generated Dox-inducible single-gene knockout cell lines and measured nuclear and cell sizes in these knockout cells. We confirmed that knockouts of most of these genes resulted in N/C ratio changes consistent with the screening results (**Fig.1i, j** and **Fig. S2**). Though, some knockouts failed to produce statistically significant changes in N/C ratio, likely due to an insufficient knockout efficiency and a reduced fitness of cells upon gene disruption (**Fig. S3** and **Supplementary Data 3**).

The presence of known regulators among the top hits in our CRISPR screen, as well as the robustness of sequencing results across 5 independent genome-wide libraries, supports the validity of our ICS-based CRISPR-based screening strategy. Overall, this enables us to systematically and confidently identify a large class of genes affecting the N/C ratio.

### Protein and RNA metabolism pathways modulate the N/C ratio by shifting the colloid osmotic pressure balance between the nucleus and the cytoplasm

Having identified genes whose disruption alters the N/C ratio, we next sought to classify these genes according to their functional roles. To this end, we performed enrichment analysis using CORUM complexes and GO biological processes (**Fig. 2a, b** and **Supplementary Data 4**). Among low N/C hits, CORUM analysis revealed a strong enrichment of complexes associated with transcriptional regulation, including the Mediator complex, RNA polymerase II-associated complexes, and MYC-MAX complexes, indicating that suppression of transcription leads to a decreased N/C ratio. GO analysis further identified enrichment of biological processes related to vesicle fusion, autophagy, chromatin organization, and mTORC1 signalling. In contrast, high N/C hits were strongly enriched for mitochondrial-related components across both CORUM and GO analyses, including mitochondrial ribosome and respiratory chain complexes, as well as processes such as mitochondrial translation and ATP synthesis. In addition, components related to chromatin regulation, cytoskeletal organization and membrane trafficking, including the Arp2/3 complex and Golgi-associated complexes, were also identified. These results suggest that regulators of the N/C ratio are associated with distinct biological processes.

**Fig. 2.**
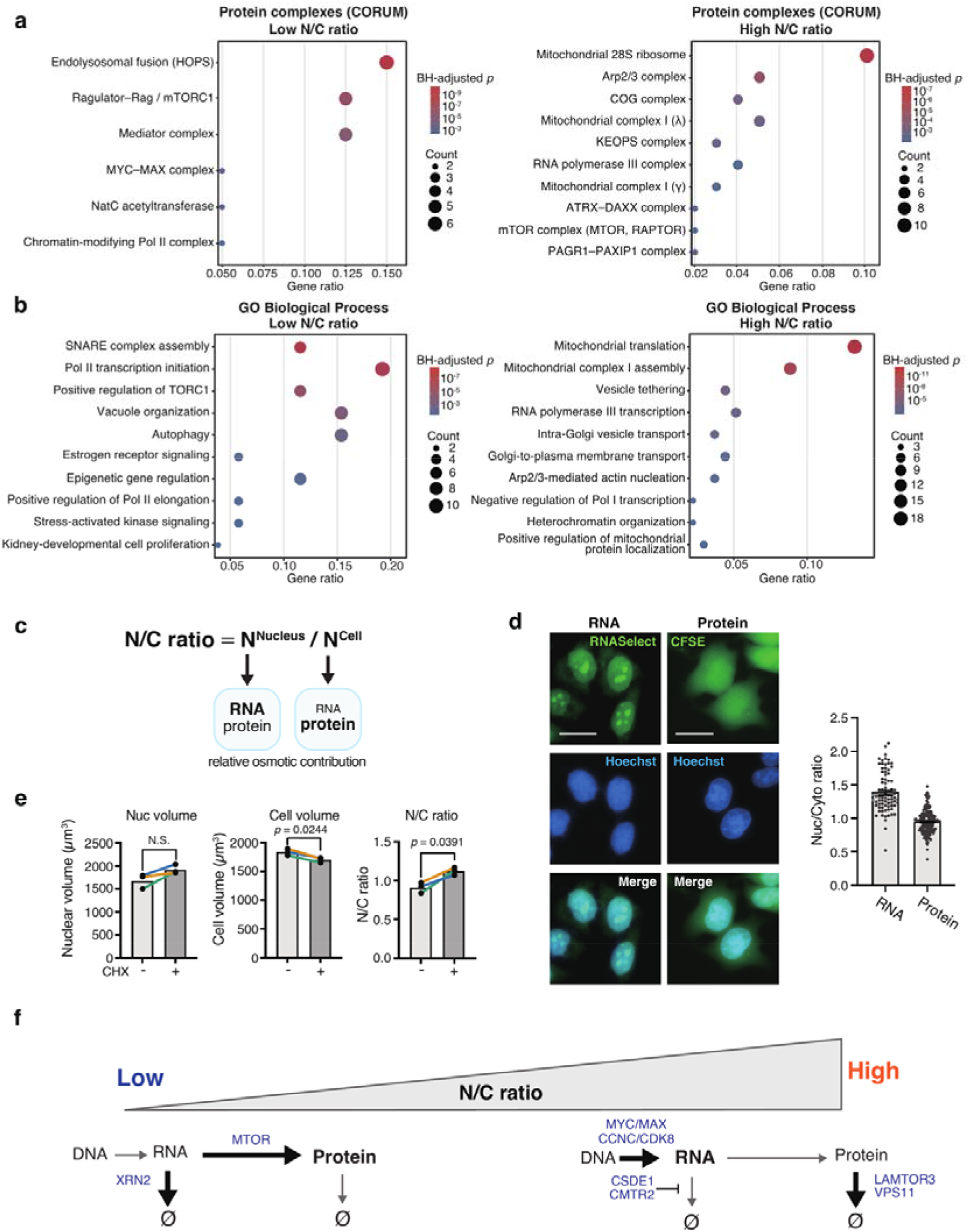
RNA and protein metabolism pathways determine the N/C ratio by setting the osmotic balance between the cytosol and the nucleoplasm. **a**, **b**, CORUM complex (**a**) and GO biological process (BP) (**b**) enrichment analysis of genes enriched in low- and high-N/C ratio cell populations. **c**, Schematic representation of the relative osmotic contributions of RNA and protein to the N/C ratio. RNA is enriched in the nucleus, whereas protein is more evenly distributed across the cell. **d**, Representative images of HeLa cells stained with RNASelect (RNA) or CFSE (protein), together with Hoechst staining to visualize nuclei. RNASelect labels total cellular RNA, while CFSE labels total cellular protein content. Merged images of RNASelect or CFSE with Hoechst are shown in the bottom row. Nuclear-to-cytoplasmic ratios of RNA and protein content are shown on the right. Each dot represents a single cell. Scale bars, 20 µm. **e**, Nuclear volume (left), cell volume (middle), and N/C ratio (right) in HeLa cells after 24 h treatment with cycloheximide (CHX). Each dot represents an independent biological replicate, and lines connect paired measurements from the same experiment. *p*-values were assessed using a two-tailed paired Student’s t-test. n = 3 biological replicates, each including at least 1,000 cells. **f**, A schematic showing how the activity of different RNA- and protein-metabolism genes across the central dogma axis can set low- or high-N/C ratio states. Examples of genes identified in the CRISPR screens are indicated in blue.

While the list of identified N/C regulator genes was functionally diverse, we observed a noticeable enrichment of genes controlling RNA and protein abundance among the top-scoring screen hits (**Fig. 1j**, **2a, b**, and **Supplementary Data 2**). We therefore hypothesized that RNA and protein metabolism may regulate the N/C ratio by altering the relative abundance of macromolecules in the nucleus and cytoplasm, thereby shifting the osmotic pressure balance between these compartments (**Fig. 2c**). This hypothesis is consistent with the colloid osmotic pressure model of nuclear size control, in which the relative abundance of macromolecules in the nucleus and cytoplasm determines the osmotic pressure balance between these two compartments, thus biophysically setting the N/C volume ratio^27–29^.

Having found that perturbations of RNA and protein metabolism genes alter the N/C ratio, we next sought to determine the intracellular distribution of these two types of macromolecules to evaluate their potential effects on the colloid osmotic pressure balance. To do this, we stained HeLa cells with fluorescent dyes labelling total cellular protein (CFSE) or RNA (RNASelect dye) (**Fig. 2d**). While protein was distributed uniformly across the cell, RNA was highly enriched in the nucleus, consistent with previous measurements in fission yeast^29^. This indicates that the RNA-to-protein ratio is substantially higher in the nucleus than in the cytoplasm, and therefore, under the colloid osmotic pressure model, one can expect that changes in the total intracellular protein-to-RNA ratio would shift the pressure balance between the nucleus and cytoplasm, and thus, alter the N/C volume ratio. For example, increased RNA abundance without a proportional increase in protein abundance would lead to a more pronounced macromolecule concentration increase in the nucleus than in the cytoplasm, and thus perturbations increasing the RNA abundance would be expected to upregulate the N/C volume ratio. Similarly, because protein is distributed broadly throughout the cell, decreasing intracellular protein abundance would be expected to reduce cell volume more than nuclear volume, thereby increasing the N/C ratio. Consistent with this prediction, cycloheximide (CHX) for 24 h significantly decreased cell volume without a significant change in nuclear volume, resulting in an overall increased N/C ratio (**Fig. 2e**).

Building on this result, we asked whether validated screen hits produced compartment-specific size changes consistent with the model. The knockout phenotypes fell into two broad phenotypic groups: some altered the N/C ratio mainly through changes in nuclear size, with little effect on cell size, whereas others acted mainly through changes in cell size, with little effect on nuclear size (**Fig. 1i, j**). We selected representative genes involved in RNA metabolism, protein synthesis, and protein turnover, and quantified their effects on nuclear and cell size. Consistent with the prediction, RNA-related perturbations primarily affected nuclear size. Knockout of the mRNA exonuclease *XRN2* primarily increased nuclear size by 1.14-fold, whereas knockout of the mRNA stability factor *CSDE1* reduced nuclear size to 0.88 of the original size (**Fig. 1i** and **Fig. S2**). In both cases, changes in cell size were minimal (1.03-fold for *XRN2* and 1.01-fold for *CSDE1*). In contrast, perturbations affecting protein homeostasis primarily affected cell size. Loss of *MTOR* led to a reduction in cell size (0.90-fold), whereas knockout of lysosomal and autophagy-related genes, including *VPS11* and *LAMTOR3*, increased cell size to 1.09-fold and 1.10-fold, respectively (**Fig. 1i** and **Fig. S2**). Notably, these perturbations had little effect on nuclear size (1.03-fold for MTOR, 1.00-fold for *VPS11*, and 1.03-fold for *LAMTOR3*). Together, these results indicate that RNA-related perturbations preferentially affect nuclear size, whereas perturbations affecting protein synthesis, lysosomal function, and autophagy preferentially affect cell size (**Fig. 1j** and **2c**). Additionally, in our GO molecular function enrichment analysis we identified multiple E3 ubiquitin ligase genes among the high-N/C-ratio hits, most of which have been reported to localize to the nucleus^37–44^ (**Fig. S4** and **Supplementary Data 4**). This observation suggests that nuclear protein turnover may also contribute to nuclear size regulation.

In addition to osmotic control on nuclear size, our experiments also revealed that certain genes involved in non-osmotic processes also influenced nuclear size: knockout of the lipid transport-related gene *ARFRP1* increased nuclear size, whereas knockout of the nuclear lamina component *LMNA* reduced nuclear size (**Fig. 1i** and **Fig. S2**). Moreover, functional enrichment analyses of CRISPR screen data revealed enrichment of cytoskeleton-related complexes, including the Arp2/3 complex, and chromatin organization/structure-related categories (**Fig. 2a, b**). These findings suggest that the N/C ratio is also regulated by structural and biomechanical inputs including lipid transport, cytoskeletal organization, nuclear lamina and chromatin organization.

Taken together, our results suggest that the balance between protein production, which mostly contributes to colloid osmotic pressure of the cytoplasm, and RNA production and maintenance, which contributes significantly to the intra-nuclear osmotic pressure, sets the osmotic balance, which controls the N/C ratio in human cells (**Fig. 2f**). However, in contrast to fission yeast *S. pombe*, which lacks conventional lamins and therefore has low nuclear membrane tension, animal cell nuclei can resist the osmotic forces through nuclear envelope tension and cytoskeleton^29^. Consistent with this notion, our data demonstrate that the osmotic mechanisms of N/C regulation are complemented in human cells by biomechanical forces, which depend on cytoskeletal regulation, nuclear lamina, lipid trafficking, and chromatin organization. Altogether these components comprise and integrated mechano-osmotic framework for N/C ratio control in animal cells.

### Mitochondrial activity affects nuclear size through altered nuclear export

In addition to RNA and protein metabolism regulators and structural genes, both CORUM and GO analyses identified multiple mitochondria-related gene categories among the top hits detected in the high N/C population, and in particular, the genes involved in mitochondrial translation and respiration (**Fig. 2a, b**). This finding was unexpected because, to our knowledge, none of these categories have previously been associated with the N/C ratio. To validate this result, we directly tested whether mitochondrial translation affects the N/C ratio. To do this, we inhibited mitochondrial translation using chloramphenicol (Cm). Consistent with the genetic screen results, treatment of HeLa cells with Cm led to a significant increase in nuclear volume, while cell volume remained largely unchanged, resulting in an increased N/C ratio (**Fig. 3a**). The same phenotype was also observed in RPE-1 cells (**Fig. S5**).

**Fig. 3.**
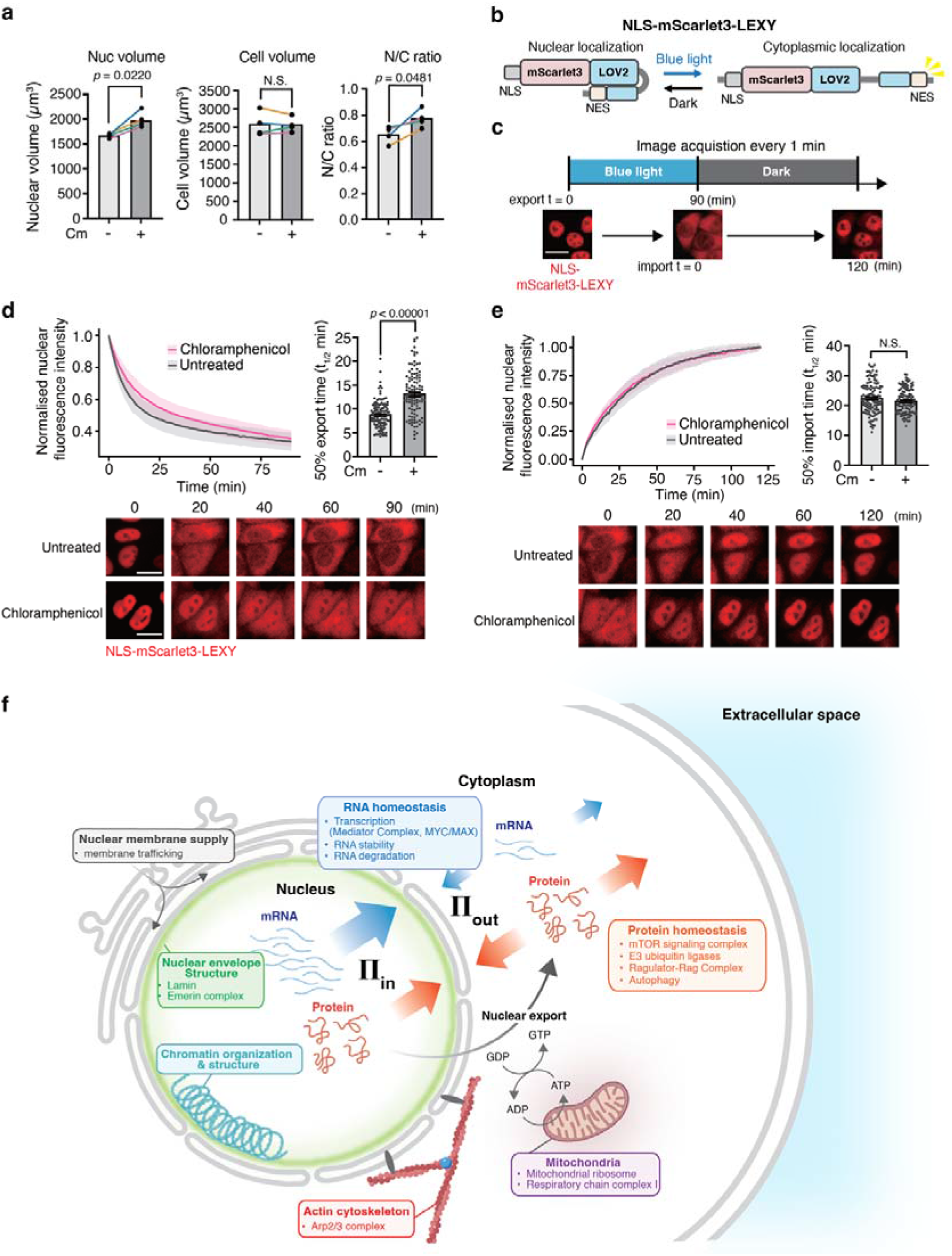
Mitochondrial translation inhibition delays nuclear export without affecting nuclear import and increases the N/C ratio. **a**, Nuclear volume (left), cell volume (middle), and N/C ratio (right) measured in HeLa cells 24 h after treatment with chloramphenicol (Cm). n = 3 biological replicates, each replicate included at least 1,000 cells. *p*-values were assessed using a two-tailed paired Student’s t-test. **b**, Schematic of the LEXY inducible protein translocation system (NLS-mScarlet3-LEXY). In the dark, mScarlet3 fluorescent protein localizes to the nucleus due to an exposed NLS, while the NES is caged within the LOV2 domain. Blue-light exposure uncages the NES, triggering nuclear export. **c**, Experimental scheme for reversible control of nucleocytoplasmic protein localization using LEXY. Cells were exposed to blue light for 90 min to induce nuclear export, followed by light withdrawal to allow nuclear import, which was monitored for 120 min. **d**, **e**, Quantification of nuclear fluorescence intensity during nuclear export (**d**) and subsequent import (**e**) using the LEXY system. Nuclear fluorescence intensity was normalized and plotted over time. Right, half-times (t_1/2_) of nuclear export (**d**) and import (**e**). Analyses were performed under untreated conditions or after chloramphenicol (Cm) treatment. Data are shown as mean ± SD (untreated: n = 122 cells; Cm: n = 111 cells). Representative images are shown below. Scale bars, 20 µm. **f**, A schematic model of the mechano-osmotic control of nuclear size in human cells. Nuclear size is governed by the balance between intra-nuclear osmotic pressure (Π_in_), driven by nuclear RNA and protein abundance, and cytoplasmic osmotic pressure (Π_out_), driven by cytoplasmic protein. RNA-related components are shown in blue, protein-related components in orange, and mitochondria in purple. Arrows indicate the proposed direction and relative strength of RNA- and protein-dependent osmotic effects acting between the nucleus and cytoplasm. The osmotic force balance is complemented by nuclear envelope biomechanics, including lamin-dependent stiffness (green) and chromatin elasticity (light blue), as well as cytoskeletal forces (red) and nuclear membrane supply (grey). Together, these factors coordinate nuclear and cell size to determine the N/C ratio.

In the context of the osmotic model, the increase in nuclear volume after Cm treatment could reflect increased retention of macromolecules within the nucleus. Because the nuclear abundance of many proteins is controlled by the balance between nuclear import and export, we next asked whether mitochondrial translation inhibition alters nucleocytoplasmic transport dynamics. To quantify nuclear export and import dynamics, we used a light-inducible nuclear export probe (NLS-mScarlet3-LEXY) (**Fig. 3b**). This probe is predominantly localized in the nucleus under dark conditions and is exported to the cytoplasm upon blue light illumination^45^. HeLa cells stably expressing this probe were used to monitor nuclear export under blue light and nuclear import after returning to dark conditions (**Fig. 3c**). Cm-treated cells delayed nuclear export (**Fig. 3d** and **Supplementary movie 1**). The half-time (t_1/2_) of nuclear export increased from 8.9 ± 0.3 min in untreated cells to 13.3 ± 0.4 min in Cm-treated cells, corresponding to an approximately 1.49-fold decrease in export rate. In contrast, the half-time of nuclear import showed no substantial change (22.6 ± 0.5 min in untreated cells versus 21.6 ± 0.4 min with Cm; **Fig. 3e**). These results indicate that inhibition of mitochondrial translation impairs nuclear export, but not nuclear import, and suggest that this may promote the accumulation of proteins within the nucleus and contribute to the observed increase in nuclear size.

Taken together, these results show that low- and high-N/C-ratio states can each arise through distinct changes in nuclear or cell size (**Fig. 3f**). Low-N/C-ratio states can result from reduced nuclear size, associated with decreased mRNA abundance or lamin A levels, or from increased cell size following impaired lysosomal function. Conversely, high-N/C-ratio states can result from increased nuclear size, associated with increased mRNA abundance, impaired vesicle trafficking, or mitochondrial dysfunction, or from reduced cell size following reduced translation or mTOR activity. Thus, diverse biological pathways regulate the N/C ratio by acting differentially on nuclear and cell size.

### Nuclear size affects human gene expression programmes independently of cell size

Having identified the regulatory pathways controlling the N/C ratio, we next sought to reveal the biological rationale for such robust and evolutionarily conserved N/C control. To address this question, we set out to identify the specific genes and pathways affected by changes in the N/C ratio in human cells. To this end, we combined the ICS technology with bulk RNA sequencing. We sorted cultured human retinal pigment epithelial (RPE-1) cells stained with the DNA dye Vybrant DyeCycle Green into “Small Nucleus” and “Large Nucleus” G1-phase cell populations and compared the transcriptomes of these two populations (**Fig. 4a**). To isolate the effects of nuclear size from the confounding cell cycle and cell size effects, we only gated G1-phase cells of similar cell size prior to sorting them into the 10% smallest and 10% largest nuclear size bins (**Fig. S6a**). After the sorting, we confirmed the difference in nuclear size using widefield microscopy (**Fig. 4b, c**). Cells sorted into the “Large Nucleus” bin had approximately 1.6-fold larger nuclei than their counterparts in the “Small Nucleus” bin, while their cell volumes, measured using the Coulter counter, were similar (**Fig. 4c**).

**Fig. 4.**
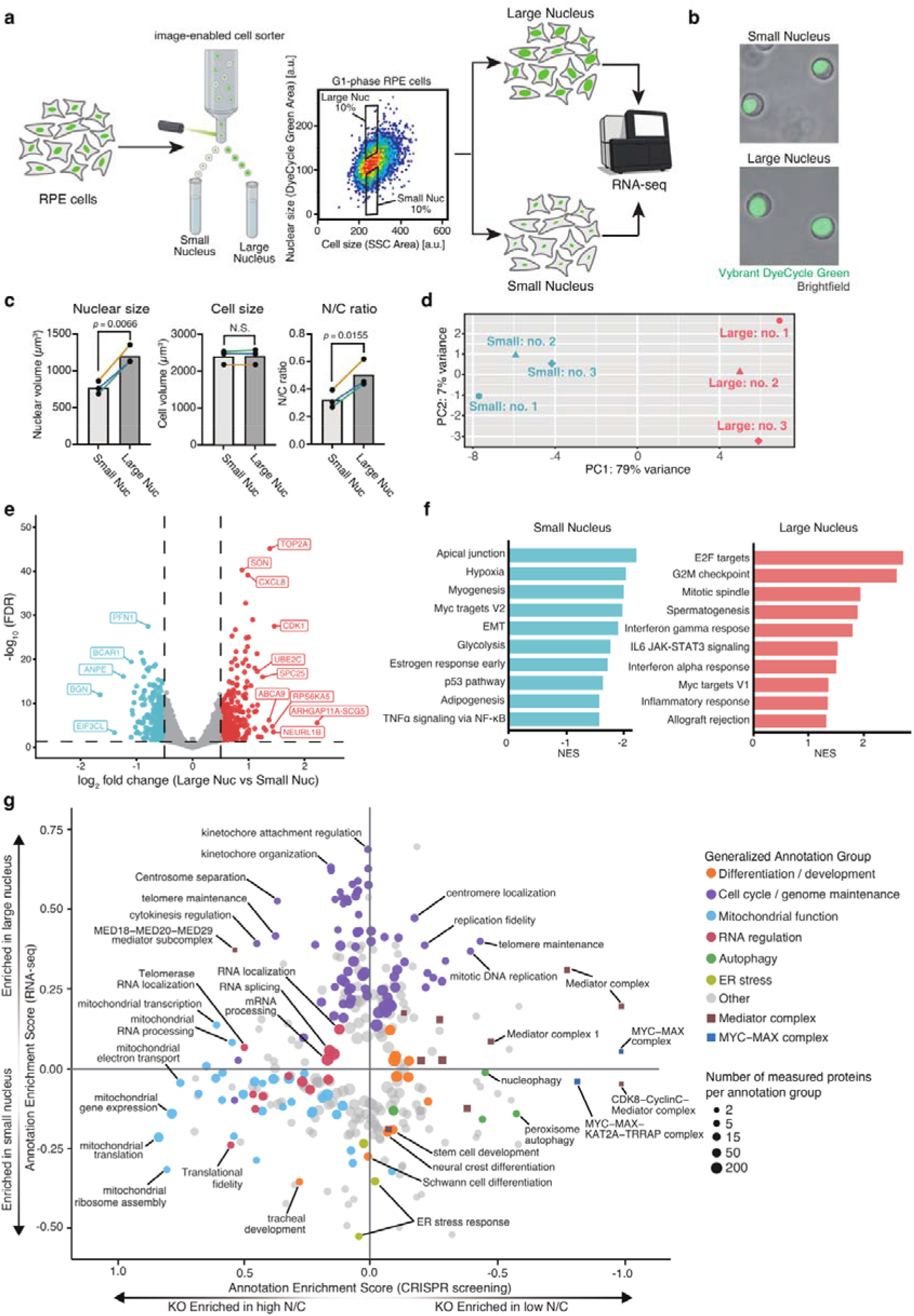
Nuclear size modulates human cell transcriptomes. **a,** Schematic overview of the experimental workflow. RPE-1 cells were stained with Vybrant DyeCycle Green and sorted using ICS. Cells of similar cell size were gated based on SSC image area, and populations comprising 10% smallest and 10% largest nuclei were identified according to nuclear size derived from fluorescence images. These populations were subsequently isolated and subjected to RNA-seq analysis. **b,** Representative widefield microscopy images of small- and large-nucleus ICS-sorted cells, showing nuclear fluorescence (Vybrant DyeCycle Green) overlaid with brightfield images. **c**, Quantification of the nuclear volume (left), cell volume (middle), and N/C ratio (right) in small- and large-nucleus ICS-sorted cell populations. Each dot represents an average of > 1,000 cells from a biological replicate (n = 3), and lines connect paired measurements from the same experiment. *p*-values were assessed using a two-tailed paired Student’s t-test. **d**, PCA of transcriptomic profiles of small- and large-nucleus populations of cells across three biological replicates (no.1–3). **e**, Volcano plot showing differentially expressed annotated protein-coding genes between small- and large-nucleus populations. Thresholds were set at FDR < 0.05 and |log_2_ fold change| ≥ 0.5. Blue dots indicate genes significantly upregulated in the small-nucleus group, whereas red dots indicate genes significantly upregulated in the large-nucleus group. **f**, Gene set enrichment analysis (GSEA) of hallmark pathways in small- and large-nucleus populations. GSEA was performed using the MSigDB Hallmark gene sets with a pre-ranked gene list based on log_2_ fold change. Pathways with FDR < 0.05 were considered significantly enriched. **g**, 2D annotation enrichment analysis comparing functional categories identified in the CRISPR screen with those enriched in the RNA-seq dataset. GO:BP annotation groups are shown as circles, using a relaxed display threshold of Benjamini–Hochberg FDR < 0.07; this panel is exploratory and inclusion does not denote statistical significance (see Methods). CORUM annotation groups for MYC–MAX and Mediator complexes are shown as filled squares and are included despite having FDR > 0.07 to facilitate comparison with Fig. 2a. Each point represents an annotation group, with point size indicating the number of genes or mapped complex subunits within each group. The position along the axes reflects the degree of enrichment in each dataset based on rank ordering. For the CRISPR screen, annotations enriched in the high-N/C knockout population are shown on the left, whereas those enriched in the low-N/C knockout population are shown on the right. Annotation groups were manually curated into generalized functional categories and coloured accordingly.

Having isolated similarly sized G1 cells with small and large nuclei, we compared the transcriptomes of these two populations using bulk RNA-seq analysis. PCA confirmed that the samples from three biological replicates were well separated according to the nuclear size (**Fig. 4d**), indicating that transcriptomes of small- and large-nucleus cells differ significantly and consistently. Subsequent analysis identified a large subset of genes differentially expressed between cells with small and large nuclei (**Fig. 4e**, **Fig. S6b**, and **Supplementary Data 5**), suggesting that nuclear size is associated with global transcriptional rearrangements that are independent of cell size and the cell cycle. Because changes in cell size were previously reported to globally impact transcriptomes and proteomes ^46–50^, we next tested whether the nuclear-size dependent transcription changes we observed in our experiment are similar to the previously reported cell size-dependent transcriptional changes. We performed a gene-matched comparison between nuclear-size-dependent transcriptional changes measured in our RPE-1 RNA-seq dataset and previously reported cell-size-dependent mRNA changes measured in the same cell typeyou^50^. We found little similarity between these two datasets (**Fig. S6c**), indicating that nuclear size affects human gene expression programmes largely independently of cell size.

To reveal functional consequences of the observed nuclear size-dependent transcriptional changes, we performed a gene set enrichment analysis (GSEA) using hallmark gene sets (**Fig. 4f** and **Supplementary Data 6**). We found that the transcriptomes of small-nucleus cells were enriched for developmental and differentiation-related programmes, including myogenesis and adipogenesis, as well as glycolysis metabolic programme and stress-responsive programmes, such as hypoxia, p53 pathway, and TNFα signalling via NF-κB. In contrast, large-nucleus cell transcriptomes were enriched for genes regulating cell cycle and mitotic programmes, including E2F targets, the G2/M checkpoint, and mitotic spindle assembly, alongside several immune and inflammatory programmes. These results suggest that nuclear size can bias cells toward distinct functional states.

While we observe significant differences in gene expression profiles between small- and large-nucleus cells, it is worth noting that formally, such gene expression differences could either be a cause or a consequence of nuclear size changes, and RNA-seq alone cannot distinguish between these two possibilities. However, the CRISPR screen, described above, provides us with a cue for determining the direction of causality, because it contains the information about genes whose expression can shift the N/C ratio. We therefore compared the pathway enrichment scores between the RNA-seq and CRISPR screen analyses (**Fig. 4g** and **Supplementary Data 7**). We reasoned that if genes from a given pathway were upregulated in small-nucleus cells according to the RNA-seq, and disruption of its genes shifted cells toward a high N/C ratio in our CRISPR screen, then the pathway was likely an upstream regulator that reduces the N/C ratio (**Fig. 4g**, lower left). Similarly, if a pathway was enriched in large-nucleus cells, and disruption of its genes shifted cells toward a low N/C ratio, the pathway was considered a candidate upstream regulator that normally increases the N/C ratio (**Fig. 4g**, upper right). Conversely, when the direction of RNA-seq enrichment did not correspond to the CRISPR phenotype, the expression difference was more likely to reflect a downstream consequence of nuclear-size variation.

When we compared the RNA-seq and CRISPR screen pathway enrichment scores, we observed only a modest overall correlation between the two datasets, indicating that most transcriptional changes detected by RNA-seq are likely consequences rather than causes of N/C ratio variation (**Fig. 4g**). RNA-seq analysis showed that, for example, development- and differentiation-related annotation terms, including stem cell development, neural crest differentiation, and Schwann cell differentiation, were positioned towards enrichment among genes upregulated in small-nucleus cells at the relaxed display threshold used for this panel (see Methods), while many cell cycle genes were upregulated in large-nucleus cells. By contrast, these terms showed no consistent directional shift in the CRISPR screen. Thus, the associated transcriptional changes are more likely to arise as consequences of nuclear size variation than to represent causal regulators of the N/C ratio. In contrast, mitochondrial annotation terms, including mitochondrial gene expression, mitochondrial translation, and mitochondrial ribosome assembly, were enriched toward the small-nucleus side of the RNA-seq analysis and strongly shifted toward the high-N/C side of the CRISPR screen. This concordance supports a role for mitochondrial gene-expression pathways as regulators that restrain the N/C ratio. We further included the Mediator and MYC–MAX complexes, two major transcriptional regulatory complexes highlighted by the CORUM analysis in **Fig. 2a**. These terms showed no consistent shift toward either small- or large-nucleus cells in the RNA-seq analysis but were shifted toward the low-N/C side of the CRISPR screen. These findings suggest that the Mediator and MYC–MAX complexes act as regulators that promote or maintain a higher N/C ratio.

Together, these results suggest that cell cycle, development and differentiation-related transcriptional changes arise downstream of altered N/C ratio, whereas mitochondrial pathways and major transcriptional regulatory complexes act upstream to regulate the N/C ratio.

### Reduced nuclear size promotes gene expression programmes associated with development and differentiation

Consistent with the Hallmark GSEA (**Fig. 4f**), GO analysis of genes expressed at higher levels in small-nucleus cells identified several developmental categories among the top-ranked terms, including positive regulation of nervous system development, positive regulation of neurogenesis, and positive regulation of cell development (**Fig. 5a** and **Supplementary Data 8**). Enrichment at the level of these individual developmental GO terms was modest (BH-adjusted *p* = 0.06–0.18), consistent with a broadly distributed transcriptional shift rather than activation of a discrete pathway. To further characterize this development- and differentiation-related transcriptional signature, we examined genes within the “stem cell development” annotation group, which was among the development-related categories positioned towards small-nucleus enrichment in **Fig. 4g** (source-specific FDR = 0.06). These genes represented distinct aspects of developmental regulation, including morphogen signalling, stem/progenitor-cell regulation, and neural crest and mesenchymal transcriptional programmes (**Fig. 5b**). The developmental regulators BMP4, HES1, and JAG1 were expressed at significantly higher levels in small-nucleus cells than in large-nucleus cells (**Fig. 5c**). This is notable because nuclear size is known to decrease during pluripotent stem cell differentiation^13–15^. We therefore hypothesized that a reduction in nuclear size during differentiation may not just be a passive result of differentiation-associated transcriptional programme activation. Instead, it might play an active role in regulating differentiation-specific gene expression programmes. To test this hypothesis and determine whether the nuclear-size-dependent transcriptional changes we observed resemble natural developmental programmes, we compared our RNA-seq results with published datasets obtained throughout stem cell differentiation trajectories. We found that gene expression changes observed in RPE-1 cells with smaller nuclei resembled those occurring during hESC differentiation into RPE-1 cells (**Fig. 5d** and **Supplementary Data 9**). This overlap was most pronounced for developmental and cell cycle genes. Similar trends were also observed when our RNA-seq results were compared with three additional stem cell differentiation datasets, including neural, endodermal, and cardiomyocyte differentiation (**Fig. S7a-c** and **Supplementary Data 9**). These results indicate that many transcriptomic changes taking place throughout the early development may occur as a downstream effect of nuclear size changes.

**Fig. 5.**
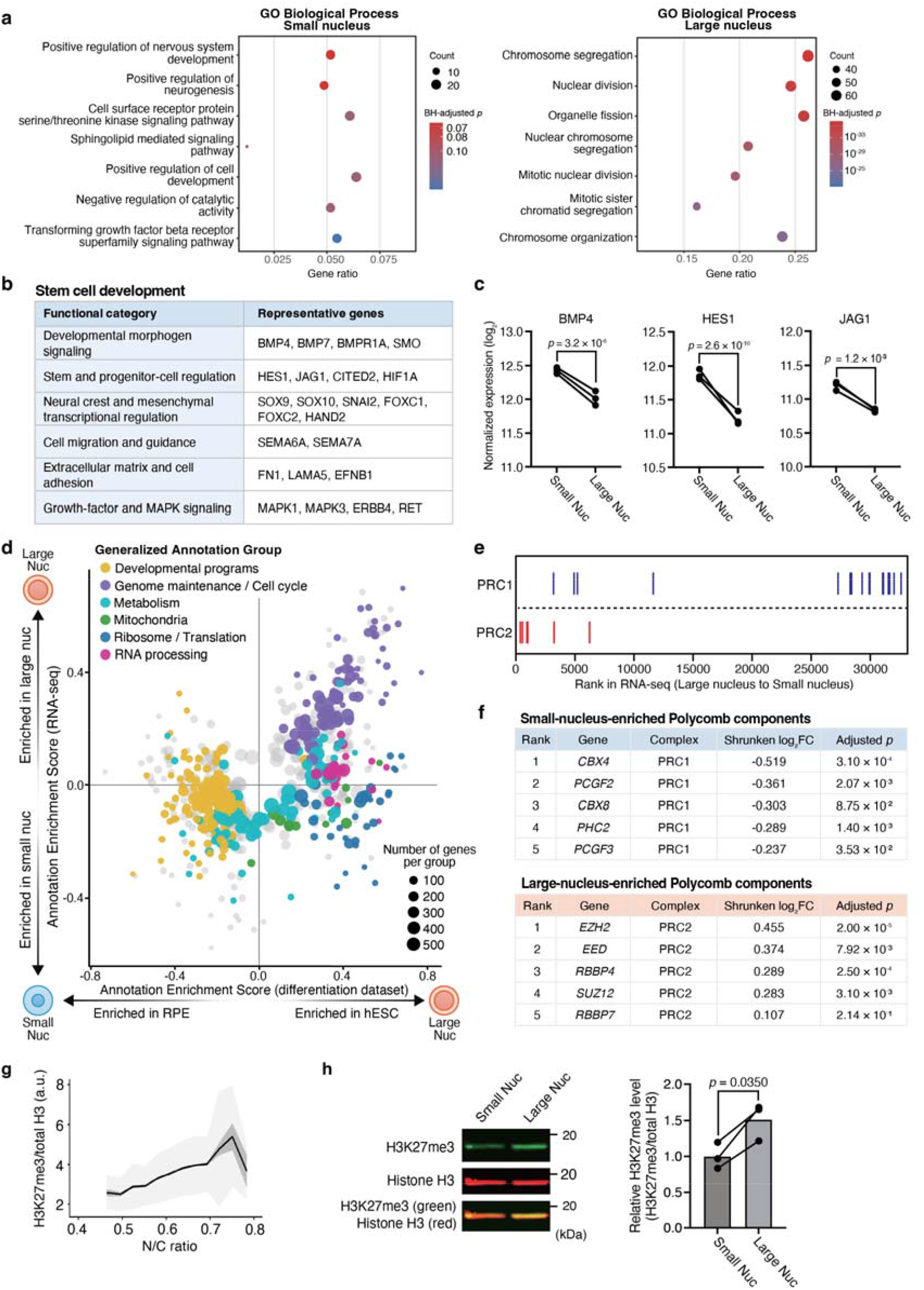
Decrease in N/C ratio promotes gene expression programmes associated with development and differentiation. **a**, GO Biological Process enrichment analysis of genes upregulated in small- and large-nucleus populations of RPE-1 cells. Dot plots show the seven highest-ranked GO terms for each group, ranked by BH-adjusted *p*-value; no significance threshold was applied, and BH-adjusted p-values are indicated by colour. The highest-ranked terms for small-nucleus cells have BH-adjusted *p* = 0.06–0.18, consistent with a broadly distributed rather than pathway-specific transcriptional shift. GeneRatio indicates the proportion of genes associated with each term and dot size represents gene count. The ten highest-ranked terms for each group are listed in **Supplementary Data 8**. **b**, **c**, Representative genes from the “stem cell development” annotation group shown in Fig. 4g that exhibited higher expression in small-nucleus cells (shrunken log_2_ fold change < 0), organized by functional category (**b**). Expression of example genes from this group (BMP4, HES1 and JAG1) in RPE-1 cells with small or large nuclei (**c**). Expression values are shown as log_2_(DESeq2-normalized counts + 1), and paired samples from the same experimental day are connected by lines (n = 3). Adjusted *p*-values were obtained from DESeq2 differential expression analysis with Benjamini–Hochberg correction. **d**, 2D annotation enrichment analysis comparing transcriptional changes between small- and large-nucleus populations with gene expression changes in differentiation datasets (RPE-1 versus hESC, GSE127352). Gene annotation groups with BH-adjusted FDR < 0.02 are shown^89^. Each dot represents an annotated gene group, with dot size indicating the number of genes in the group. The position along each axis reflects the degree of coordinated change in both datasets based on rank ordering. Gene groups were manually curated into broader annotation categories and coloured accordingly. **e, f**, Distribution of PRC1 and PRC2 component genes across the RNA-seq gene list ranked by shrunken log_2_ fold change, from large-nucleus-enriched to small-nucleus-enriched genes. Each vertical line represents an individual PRC1 (blue) or PRC2 (red) component gene (**e**). Top-ranked Polycomb component genes at the small-nucleus-enriched (upper) and large-nucleus-enriched (lower) ends of the ranked gene list, together with their shrunken log_2_ fold changes and adjusted *p*-values (**f**). **g**, Immunofluorescence analysis of H3K27me3 across N/C ratios using an imaging-enabled cell sorter. The N/C ratio is calculated from measured nuclear and cell areas. H3K27me3 levels were normalized to total histone H3. Cells were binned by the N/C ratio, black line indicates bin means, dark shading indicates SEM, and light shading indicates SD for approximately 100,000 cells. **h**, Immunoblotting of H3K27me3 and total histone H3 in sorted small- and large-nucleus RPE-1 populations. Left – a representative immunoblot image, right – quantification. Total histone H3 was used for normalization of H3K27me3. Each dot represents an individual biological replicate (n = 3), and lines connect paired measurements from the same experiment. *p*-values were assessed using a two-tailed paired Student’s t-test.

To identify potential regulatory mechanisms underlying nuclear size-dependent transcriptional changes, we focused on the most differentially expressed genes between small- and large-nucleus cells. Among the genes with strongest nuclear-size dependence we observed components of the Polycomb repressive complexes PRC1 and PRC2, which changed in opposite directions. Most components of the canonical PRC1 were enriched in small-nucleus cells, whereas PRC2 core components were enriched in large-nucleus cells (**Fig. 5e, f**). Polycomb complexes are key epigenetic regulators that mediate gene repression and play a central role throughout embryo development.

During early stages of embryonic development, PRC2 complex is essential for maintaining self-renewal and pluripotency of embryonic stem cells, while keeping them in a “poised” state. PRC2 represses developmental genes by catalysing H3K27 trimethylation (H3K27me3) and thus placing a repressive epigenetic mark on them, which leads to heterochromatin formation and lineage-specific gene silencing^51,52^. As pluripotent cells undergo lineage specification during later stages of embryo development, the activity of PRC2 partially decreases, while canonical PRC1 starts playing the key role in controlling differentiation programmes^53,54^. PRC1 can be recruited to H3K27me3 sites to maintain chromatin compaction^55^ and stable silencing of lineage-specific developmental genes to ensure accurate cell fate specification^56,57^. This is consistent with our observation that PRC2 is enriched in cells with larger nuclei, which are morphologically more similar to pluripotent stem cells, while canonical PRC1 is enriched in smaller-nuclei cells, which are more similar to the cells undergoing lineage specification.

Having found that PRC2 component mRNAs were more highly expressed in large-nucleus cells, we examined whether this increase was accompanied by elevated H3K27me3 levels – a functional output of PRC2 activity. Immunofluorescence-based flow cytometry showed that H3K27me3 levels increased with increasing N/C ratio (**Fig. 5g**). Consistent with this result, immunoblotting of sorted small- and large-nucleus populations showed higher H3K27me3 levels in large-nucleus cells than in small-nucleus cells (**Fig. 5h**).

Together, these findings suggest that decreasing N/C ratio is associated with a transition from a PRC2-dominant to a PRC1-dominant Polycomb state. This transition is accompanied by reduced H3K27me3, consistent with the differentiation-like transcriptional programmes observed in small-nucleus cells. Thus, changes in nuclear size likely play an active role in the reorganization of Polycomb-associated epigenetic states during differentiation.

### Small-nucleus mESCs more readily exit pluripotency and initiate neural differentiation in response to retinoic acid

Having found that a decrease in nuclear size promotes gene expression changes resembling those observed during pluripotent stem cell differentiation, we hypothesized that such transcriptome rearrangements may bias stem cells with smaller nuclei towards differentiation programmes. To test this hypothesis and determine whether smaller nuclear size promotes differentiation in mouse embryonic stem cells (mESCs), we used the ICS approach to separate G1-phase mESCs of similar cell size into small- and large-nucleus populations, re-plated them and exposed to a differentiation protocol (**Fig. 6a** and **Fig. S8**). Like before, we quantified nuclear volume after sorting by widefield microscopy, while cell volume was measured using the Coulter counter (**Fig. 6b, c**). The average nuclear volume of the large-nucleus population was approximately 1.81-fold greater than that of the small-nucleus population, whereas cell volume did not differ significantly between the two populations. Consequently, the two populations had clearly distinct N/C ratios.

**Fig. 6.**
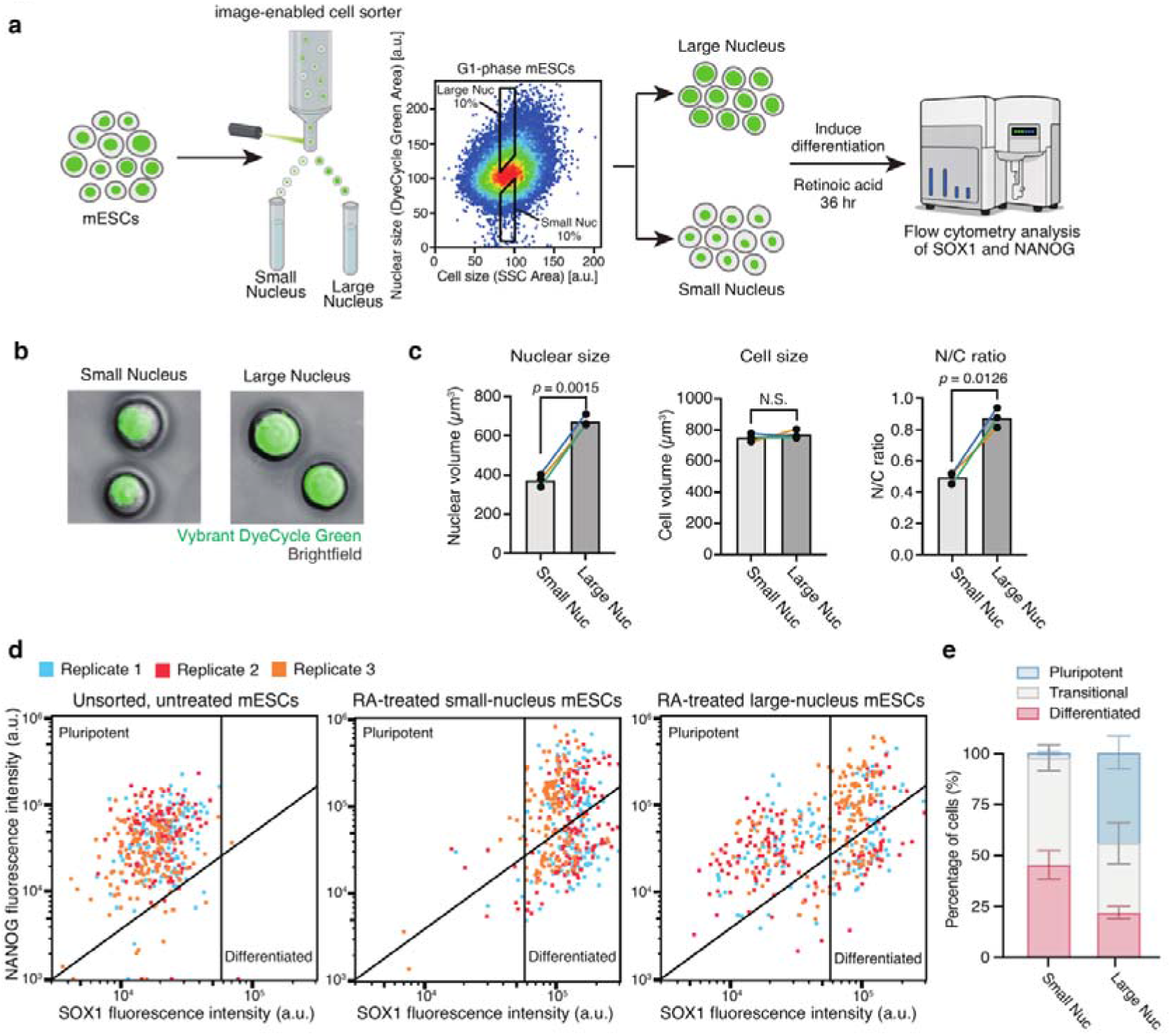
mESCs with smaller nuclei are more prone to exiting pluripotency and initiating differentiation upon retinoic acid stimulation. **a,** Schematic overview of the experimental workflow. mESCs were stained with Vybrant DyeCycle Green and sorted using the ICS. G1-phase cells of similar cell size were gated based on SSC, and the populations comprising the 10% smallest and 10% largest nuclei were isolated based on Vybrant DyeCycle Green staining area. The sorted cells were treated with RA for 36 h and subsequently analysed for SOX1 and NANOG expression by flow cytometry. The complete gating strategy is shown in Fig. S8. **b,** Representative widefield microscopy images of sorted small- and large-nucleus mESCs, showing nuclear fluorescence from Vybrant DyeCycle Green overlaid with brightfield images. **c,** Quantification of the nuclear volume (left), cell volume (middle), and N/C ratio (right) in small- and large-nucleus populations. Each dot represents an average of > 1,000 cells from an individual biological replicate (n = 3), and lines connect paired measurements from the same experiment. *p*-values were assessed using a two-tailed paired Student’s t-test. **d,** SOX1 and NANOG expression distributions in unsorted untreated mESCs and in RA-treated small- and large-nucleus mESC populations. For visualization, 140 single-cell events were downsampled from each of three independent experiments and overlaid, with each experiment shown in a different colour. SOX1□/NANOG□ cells were classified as pluripotent, whereas SOX1□/NANOG□ cells were classified as differentiated. **e,** Quantification of pluripotent (SOX1□/NANOG□), differentiated (SOX1□/NANOG□), and transitional subpopulations in the ICS-sorted small- and large-nucleus mESC populations after 36 h of RA treatment. The transitional category comprises both SOX1□/NANOG□ and SOX1□/NANOG□ cells. Data are from n = 3 independent biological replicates; error bars denote SEM.

The sorted cells were then re-plated and treated for 36 h with retinoic acid (RA), which induces exit from pluripotency and promotes early neural differentiation in mESCs. After RA treatment, we harvested the cells and analyzed the expression of a pluripotency marker NANOG and early neural differentiation marker SOX1 by flow cytometry. We compared the SOX1–NANOG expression distributions of untreated mESCs, RA-treated small-nucleus mESCs, and RA-treated large-nucleus mESCs (**Fig. 6d**). Nearly all untreated mESCs were located within the SOX1□/NANOG□ region of the SOX1–NANOG scatter plot, confirming that the starting cell population was predominantly pluripotent. Following RA treatment, the proportion of SOX1□/NANOG□ pluripotent cells was significantly lower in the small-nucleus population than in the large-nucleus population – approximately 3% against 44%, respectively. At the same time, the proportion of SOX1□/NANOG□ differentiated cells was higher in the small-nucleus population – approximately 45%, compared with 22% in large-nucleus population (**Fig. 6e**).

Together, these findings reveal that nuclear size, independently of cell size, predicts differentiation competence in mESCs. Small-nucleus cells preferentially exit pluripotency and initiate neural differentiation in response to RA, whereas large-nucleus cells preferentially retain the pluripotent state. This is consistent with the nuclear size-dependent gene expression and H3K27 tri-methylation changes we described above.

## Discussion

Across diverse organisms and cell types, nuclear size scales with cell size to maintain a characteristic nuclear-to-cell volume ratio^58^, yet the mechanisms that establish this relationship and its consequences for cell physiology have remained poorly understood. Previous observations that N/C ratio changes during major cell-state transitions, including stem cell differentiation, development, and tumor progression suggested that nuclear size may be coupled to gene regulation, but whether changes in nuclear size actively influence transcriptional programmes, and how cells tune nuclear size relative to cell size remained unknown for many decades^16,58,59^.

In this study, we have demonstrated that changes in nuclear size lead to a broad range of changes in gene expression programmes. In particular, cells with smaller nuclei showed transcriptional shifts similar to those observed across multiple lineage differentiation programmes. This finding is consistent with previous observations that pluripotent stem cells generally exhibit high N/C ratios^11,12^, and that nuclear size decreases during differentiation^13–15^. To identify potential regulatory mechanisms underlying these nuclear size-dependent transcriptional changes, we examined Polycomb-associated chromatin regulation. PRC1 and PRC2 are key epigenetic regulators of cell fate transitions during development^60,61^. Cells with larger nuclei showed higher expression of mRNAs encoding PRC2 components, whereas cells with smaller nuclei showed higher expression of mRNAs encoding PRC1 components. Consistent with the increase in mRNA expression of PRC2 components, H3K27me3 levels were also elevated in large-nucleus cells. These results suggest that a shift from a PRC2-dominant state to a PRC1-dominant state may occur as the N/C ratio decreases and indicate that changes in nuclear size are associated with Polycomb-dependent chromatin regulatory states. Such a change in Polycomb state is consistent with the differentiation-associated transcriptional programmes observed in small-nucleus cells, supporting the possibility that nuclear size changes are linked to the reorganization of epigenetic states during differentiation. However, further work will be required to mechanistically determine how nuclear size regulates the expression and activity of PRC1/2 components.

Consistent with the observed gene expression changes, differences in nuclear size were also associated with distinct responses to differentiation cues. We separated G1-phase mESCs of similar cell size into small- and large-nucleus populations and induced early neural differentiation with retinoic acid. Small-nucleus mESCs more readily exited the NANOG-positive pluripotent state and entered the SOX1-positive early neural differentiation state, whereas large-nucleus mESCs were more likely to retain pluripotency following retinoic acid treatment. These findings extend the transcriptional and Polycomb-associated chromatin observations by showing that nuclear size differences present before differentiation are associated with different propensities to exit pluripotency and initiate neural differentiation in response to the same differentiation stimulus. Thus, the reduction in nuclear size that accompanies stem-cell differentiation may not be just a passive morphological consequence, but may actively promote differentiation-associated transcriptional changes and the propensity of cells to exit pluripotency. This interpretation is consistent with recent work showing that changes in the physical state of the nucleus alter chromatin organization and influence pluripotent stem-cell differentiation^62^, as well as with concurrent work showing that altering nuclear size can change the physical properties of the nucleoplasm and nuclear organization in *Schizosaccharomyces pombe*^17^ and reshape chromatin organization and transcription in *Saccharomyces cerevisiae* (*Ohsawa et al., bioRxiv 2026*).

However, two limitations should be stated. First, because small- and large-nucleus mESC populations were isolated by sorting rather than by experimentally manipulating nuclear size, we cannot exclude that nuclear size covaries with unmeasured features of cellular state — growth history, metabolic activity, or position within G1 — that independently influence differentiation propensity. Matching for cell volume and cell-cycle phase reduces the potential contribution of these two variables but does not exclude other covariates. Second, although our experiments establish an association between nuclear size and Polycomb state and show that nuclear size predicts the response to differentiation cues, they do not establish how nuclear size is sensed or transmitted to the transcriptional and epigenetic machinery. Directly imposing nuclear size before differentiation will provide a direct test of an instructive role, and the light-inducible control of nucleocytoplasmic partitioning and genetic manipulations established here (**Fig. 3b–e**) offer two possible routes.

To address the long-standing question of what sets the N/C ratio and systematically identify the molecular mechanisms controlling nuclear size, we combined high-speed image-enabled cell sorting technology with genome-wide CRISPR screening, treating the N/C ratio as a direct cellular phenotype. Our analysis demonstrated that the N/C ratio is not controlled by a single pathway, but is instead determined by the interplay of several cellular processes (**Fig. 3f**). The identified factors spanned multiple functional categories that can influence nuclear and cell size, including RNA homeostasis, protein homeostasis, mitochondrial function, nuclear membrane supply, nuclear envelope structure, chromatin organization, and the actin cytoskeleton. Targeted validation further showed that these factors do not all alter the N/C ratio in a uniform manner: some perturbations primarily affected nuclear size, whereas others mainly affected cell size. Thus, the N/C ratio is unlikely to be the outcome of a single coordinated growth mechanism, but rather reflects the combined effects of multiple pathways acting on nuclear and cell size.

Our observations are consistent with previously proposed colloid-osmotic models of nuclear size control^27–29^ and with two concurrent studies showing that changes in the nucleocytoplasmic partitioning of a massively overexpressed protein can alter the N/C ratio in budding yeast (*Ohsawa et al., bioRxiv 2026*) and fission yeast^17^. Our findings extend this framework by showing that in mammalian cells osmotic regulation works together with structural and mechanical constraints to control nuclear size. In this framework, macromolecules accumulated within the nucleus generate an internal osmotic pressure (Π_in_), which is balanced against the macromolecular content of the cytoplasm and cytoplasmic pressure (Π_out_). Our measurements showed that RNA was enriched in the nucleus, whereas protein was distributed relatively uniformly between the nucleus and cytoplasm. Therefore, changes in total cellular RNA abundance are expected to be reflected primarily in nuclear macromolecular content and thereby affect nuclear size, whereas changes in total cellular protein abundance are expected to be reflected similarly in both the nucleus and cytoplasm and primarily affect cell size (**Fig. 2c**).

Consistent with the RNA–protein osmotic balance model, pathways involved in RNA and protein metabolism emerged as major functional groups contributing to N/C ratio regulation in our screen. For RNA homeostasis, our screen identified genes associated with mRNA transcription, stabilization, and degradation. Disruption of either the mRNA exonuclease *XRN2* or the mRNA stability factor *CSDE1* altered nuclear size, with little effect on cell size. These observations are consistent with a previous genetic screen in fission yeast, which similarly identified RNA metabolism-related pathways as key regulators of nuclear size^26^. In contrast, for protein homeostasis, our CRISPR screen showed enrichment of genes associated with mTOR signalling, the lysosome-associated Ragulator–Rag complex, autophagy, and ubiquitin-mediated protein degradation. Disruption of *MTOR* reduced cell size, whereas disruption of the lysosome-related factors *LAMTOR3* and *VPS11* increased cell size; all three perturbations had little effect on nuclear size. Disruption of *POLR3H* and *ATRX* also led to reduced cell size. *POLR3H* is involved in RNA polymerase III–mediated transcription of small RNAs such as 5s rRNAs and tRNAs^63^, whereas *ATRX* has been implicated in chromatin regulation at ribosomal DNA loci^64^. The reduction in cell size following disruption of these genes is therefore consistent with a reduced capacity for ribosome biogenesis and protein synthesis. Together, these findings indicate that pathways involved in protein synthesis and degradation, ribosome biogenesis, and lysosome-associated signalling and trafficking regulate the N/C ratio primarily through changes in cell size.

Protein homeostasis may also affect nuclear size through selective changes in nuclear protein abundance. Several E3 ubiquitin ligases were found among high-N/C-ratio hits in our CRISPR screen, and most of these ligases have been reported to localize to the nucleus^37–44^. Loss of nuclear E3 ligase activity could stabilize nuclear protein substrates, thereby increasing nuclear protein abundance. Such a nuclear-specific increase in protein abundance may raise nuclear osmotic pressure and thereby increase nuclear size and the N/C ratio.

An important question for the RNA–protein osmotic balance model is whether RNA, which is substantially less abundant than protein, can make a significant contribution to nucleocytoplasmic osmotic balance. In HeLa cells, total protein content has been estimated at approximately 156 pg per cell^65^, whereas total RNA content is approximately 23 pg per cell^66^, indicating that protein is approximately sevenfold more abundant than RNA by mass. The difference is even larger when molecular abundance is considered: HeLa cells have been estimated to contain approximately 2.0–2.3×10^9^ polypeptide molecules^65^, whereas a typical mammalian cell contains roughly 10^7^ RNA molecules^67^. Thus, in terms of total molecular abundance, proteins greatly outnumber RNA molecules. However, this difference in total molecular abundance cannot be directly equated with a corresponding difference in their contributions to colloid osmotic pressure across the nuclear envelope. In the model proposed by Lemière et al.^29^, nuclear size and the N/C ratio are governed by colloid osmotic pressure generated by macromolecules in the nucleoplasm and cytoplasm. In this context, the relevant quantity is not simply the total number of macromolecules in the cell, but the number of particles that can maintain an effective concentration difference across the nuclear envelope^29^. From this perspective, not all polypeptide molecules can be counted as independent osmotically active particles. Proteome-wide quantitative mass-spectrometry analyses have shown that highly abundant proteins tend to be relatively small, with proteins in the highest-abundance range having an average molecular mass of approximately 40 kDa^68^. Proteins below approximately 40–60 kDa can undergo appreciable passive diffusion through nuclear pore complexes, with permeability decreasing as molecular size increases^69^. Thus, a substantial fraction of abundant soluble proteins may equilibrate between the nucleus and cytoplasm, limiting their ability to sustain a persistent osmotic pressure difference across the nuclear envelope. Proteins embedded in cellular membranes or sequestered within membrane-bound organelles also do not directly belong to the soluble macromolecular pools of the nucleoplasm and cytoplasm. Furthermore, when multiple polypeptides are assembled into a large macromolecular complex, the individual polypeptide subunits do not behave as independent osmotic particles^70^. Thus, the effective number of protein particles contributing to the osmotic pressure difference across the nuclear envelope may be substantially smaller than the total number of polypeptide molecules. A similar distinction applies to RNA: the total number of RNA molecules cannot be directly equated with the number of RNA-containing particles that contribute to an osmotic pressure difference across the nuclear envelope. Unlike many relatively small soluble proteins, many RNAs, including mRNAs and rRNAs, associate with proteins to form large RNP complexes that are not expected to freely equilibrate across nuclear pore complexes by passive diffusion and instead undergo regulated nuclear export^71^. Consequently, at least some nuclear RNA-containing particles may not rapidly equilibrate between the two compartments and could therefore maintain a concentration difference across the nuclear envelope. Thus, although proteins greatly outnumber RNAs in total molecular abundance, this ratio cannot be assumed to directly reflect their relative contributions to the osmotic pressure difference across the nuclear envelope. Consistent with this distinction, an accompanying study by *Ohsawa et al.* estimates that approximately 7.2 million osmotically active colloid particles determine the N/C ratio in budding yeast, roughly one tenth of the ∼75 million protein molecules inferred from proteomics. The effective number of osmotically active protein particles is therefore likely far lower than the total number of protein molecules, making the contribution of RNA greater than abundance alone suggests. Our genetic data bear on this directly: loss of *XRN2* or *CSDE1* altered nuclear volume with little effect on cell volume, whereas perturbing protein synthesis or degradation did the converse.

Notably, genes associated with mitochondrial function – in particular, mitochondrial translation and respiratory chain complex I – were strongly enriched among the high-N/C-ratio hits in our screen, which was unexpected. Consistent with this result, chloramphenicol-mediated inhibition of mitochondrial translation increased nuclear size without substantially altering cell size. By measuring the nuclear export and import dynamics of the light-inducible nuclear export LEXY construct, we found that inhibition of mitochondrial translation delayed NES-dependent nuclear export, while having little effect on nuclear import, suggesting that mitochondrial function may regulate the N/C ratio through nucleocytoplasmic redistribution of osmotically active macromolecules. This result is consistent with published observations that NES-dependent nuclear export requires RanGTP-dependent export complex formation and is more sensitive to reduced RanGTP availability than classical NLS-dependent nuclear import^72–74^. Thus, inhibition of mitochondrial function may selectively impair Ran-dependent nuclear export, likely through changes in cellular energy state, thereby promoting the accumulation of proteins in the nucleus. This shift in nucleocytoplasmic protein distribution may contribute to nuclear enlargement by increasing nuclear osmotic pressure.

Apart from energy-dependent effects of mitochondria on nucleocytoplasmic transport, another possibility is that disruption of mitochondrial proteostasis, caused by mitochondrial ribosome gene knockout or chloramphenicol treatment, activates compensatory mitochondrial retrograde signaling via the mitochondrial unfolded protein response (UPR^mt^) and integrated stress response (ISR) pathways, which in turn upregulate transcription of nuclear-encoded mitochondrial genes^75–79^. Such upregulation could then shift the protein-to-RNA osmotic balance between the cytoplasm and the nucleus, because the upregulated transcripts are produced in the nucleus, whereas the proteins they encode are imported into mitochondria and thus, sequestered from the cytoplasm. It is also plausible that mitochondrial activity may intersect with the structural and mechanical arm of nuclear-size control through metabolite-dependent regulation of chromatin. Mitochondrial dysfunction resulting in reduced acetyl-CoA availability can affect histone acetylation^80,81^, whereas histone-modification-dependent changes in chromatin compaction can alter nuclear mechanics and morphology^32^. Thus, mitochondrial activity may influence nuclear size through both osmotic and chromatin-mechanical routes, and further testing and delineation of these pathways will be required to gain a mechanistic understanding of how mitochondrial function regulates the N/C ratio.

Despite the evident importance of colloid osmotic pressure in nuclear and cell size regulation, the N/C ratio in human cells cannot be explained solely by the osmotic effects arising from RNA and protein abundance and nucleocytoplasmic distribution. While this model may be sufficient for fission yeast cells, which lack lamins and have a negligible nuclear membrane tension, additional structural and mechanical factors are likely to contribute to N/C ratio regulation in human cells^29^. In addition to pathways involved in RNA and protein abundance and intracellular distribution, our genetic screen also identified pathways related to nuclear membrane supply (membrane trafficking), nuclear envelope structure (Lamin and Emerin complex), chromatin organization, and the actin cytoskeleton (Arp2/3 complex). These pathways correspond to mechanisms previously implicated individually in nuclear size and morphology, including membrane supply to the nuclear envelope, mechanical stiffness of the nuclear lamina, physical properties of chromatin, and cytoskeletal forces^21–23,82^.

Taken together, our findings suggest that osmotic, structural, and mechanical factors act in concert to establish integrated mechano-osmotic control of the N/C ratio in human cells. In this integrated mechano-osmotic model, osmotic effects arising from RNA and protein abundance and distribution act together with membrane availability and the mechanical properties of the nuclear envelope, chromatin, and cytoskeleton to determine nuclear size relative to cell size.

An increased N/C ratio is a hallmark of cancer cells and is associated with malignant properties including altered cell proliferation, migration, invasiveness, and metastasis^83^. Identifying the molecular pathways that regulate the N/C ratio may therefore provide new opportunities to investigate how nuclear size contributes to malignant cell behaviour and, potentially, to explore whether manipulating nuclear size can influence cancer cell invasion and metastatic potential.

Taken together, our findings address a fundamental problem in cell biology that has remained unresolved for more than a century, establishing nuclear size as a physical regulator of cell state and providing a mechanistic connection between cellular architecture, gene regulation and cell fate. By systematically identifying molecular pathways that regulate nuclear size relative to cell size, we provide a means to experimentally manipulate the N/C ratio and thereby investigate its functional roles in cellular physiology. Our findings further suggest that nuclear size can influence chromatin organization, gene expression and cell fate, raising the possibility that the N/C ratio itself could be manipulated to influence cell-state transitions. In the longer term, this framework opens new opportunities to investigate how cellular architecture controls cell state and, potentially, to manipulate cell-state transitions with practical implications for regenerative biology, including the maintenance of stem-cell pluripotency, cellular reprogramming and directed differentiation.

## Figures and legends

**Figure S1.**
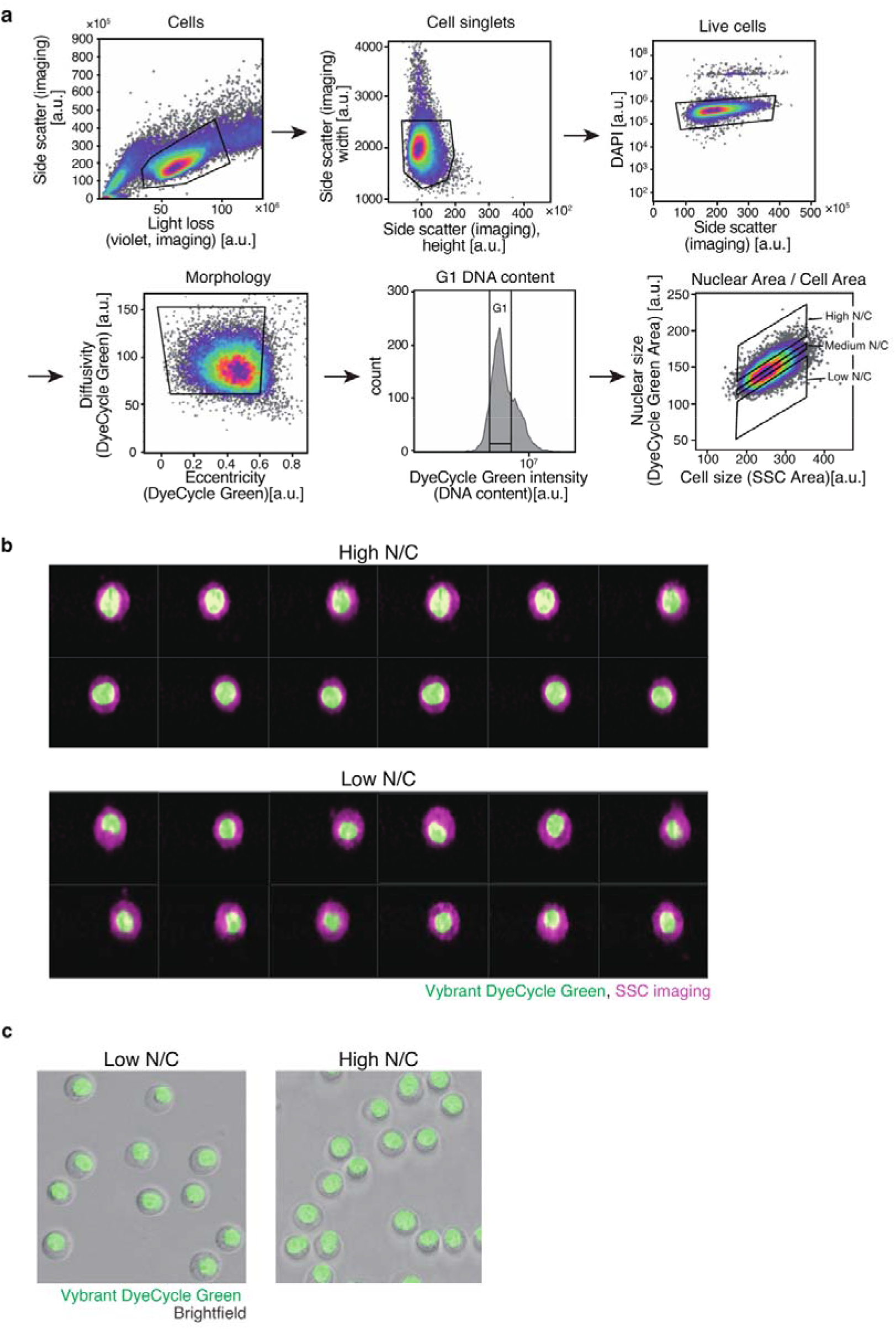
Gating strategy used in the ICS-based genome-wide CRISPR screen identifying regulators of the N/C ratio. **a**, Dox-induced Tet::Cas9 HeLa cells were sequentially gated by Light Loss vs Size Scatter to exclude debris, then by Side Scatter Height vs Side Scatter Width to select singlets, then by lack of DAPI signal to select live cells. Live cells with normal nuclear morphology were further selected based on imaging-derived shape parameters (Vybrant DyeCycle Green DNA stain eccentricity and diffusivity). G1-phase cells were identified by DNA content (Vybrant DyeCycle Green intensity). Nuclear size (Vybrant DyeCycle Green Image Area) and cell size (SSC) were quantified in the gated population to calculate the N/C ratio used for screening. 10-12% of cells with the highest, lowest and medium N/C ratios were sorted for subsequent sgRNA sequencing and enrichment analysis. **b**, Representative images from the Discover S8 cytometer, obtained using the BD CellView™ Image Technology, show cells from the collected populations, with nuclei visualized by Vybrant DyeCycle Green fluorescence (green) and cell morphology visualized by side-scatter (SSC) imaging (magenta). **c**, Representative widefield microscopy images of small- and large-nucleus populations, showing nuclear fluorescence (Vybrant DyeCycle Green) overlaid with brightfield images.

**Figure S2.**
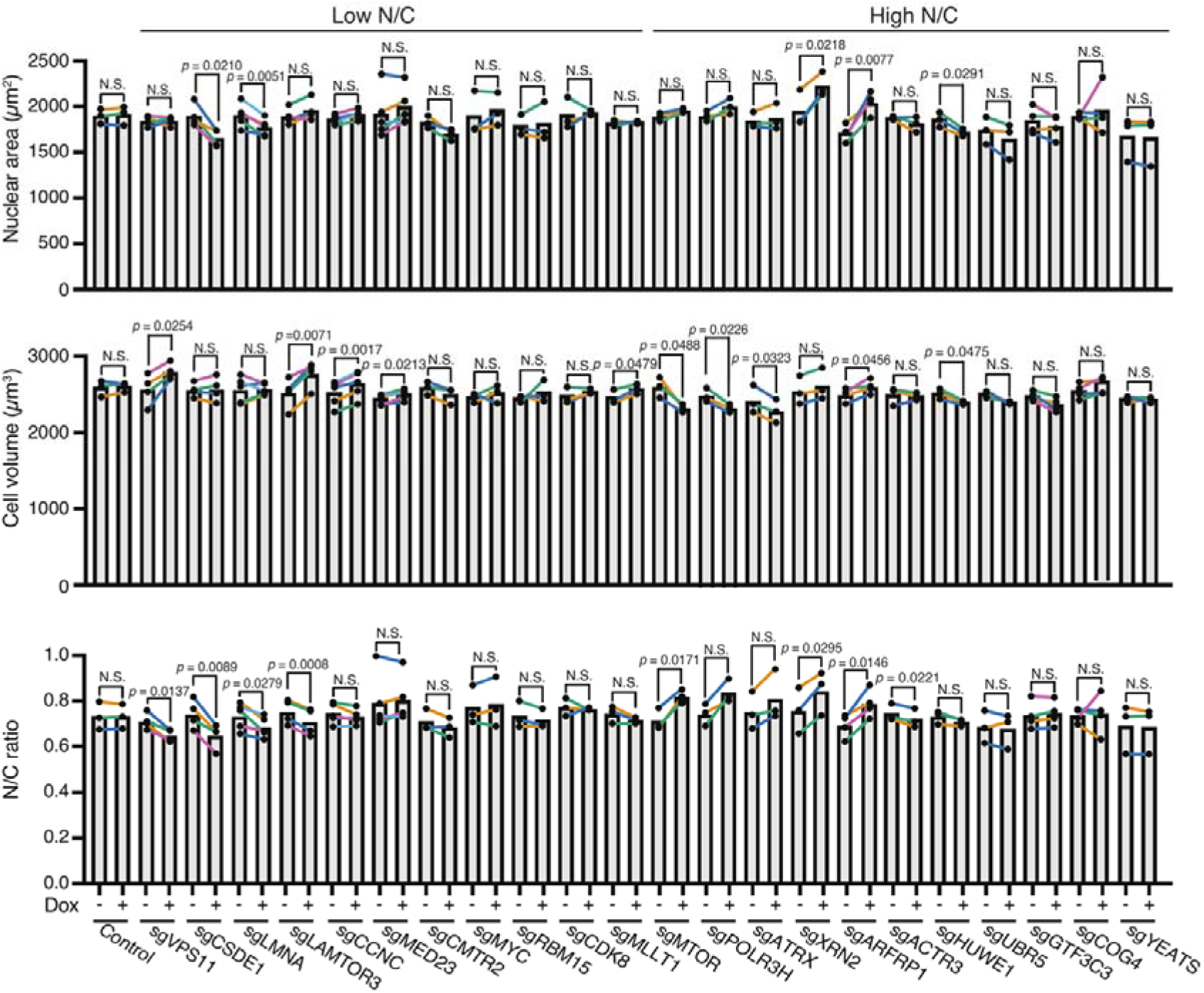
Validation of candidate genes affecting the N/C ratio. Bar plots show nuclear volume (top), cell volume (middle), and N/C ratio (bottom) for the indicated candidate genes. Tet::Cas9 HeLa cells expressing each sgRNA were analysed under untreated and Dox-treated conditions. n = 3 biological replicates. Indicated *p*-values are calculated using a paired two-tailed Student’s t-test.

**Fig. S3.**
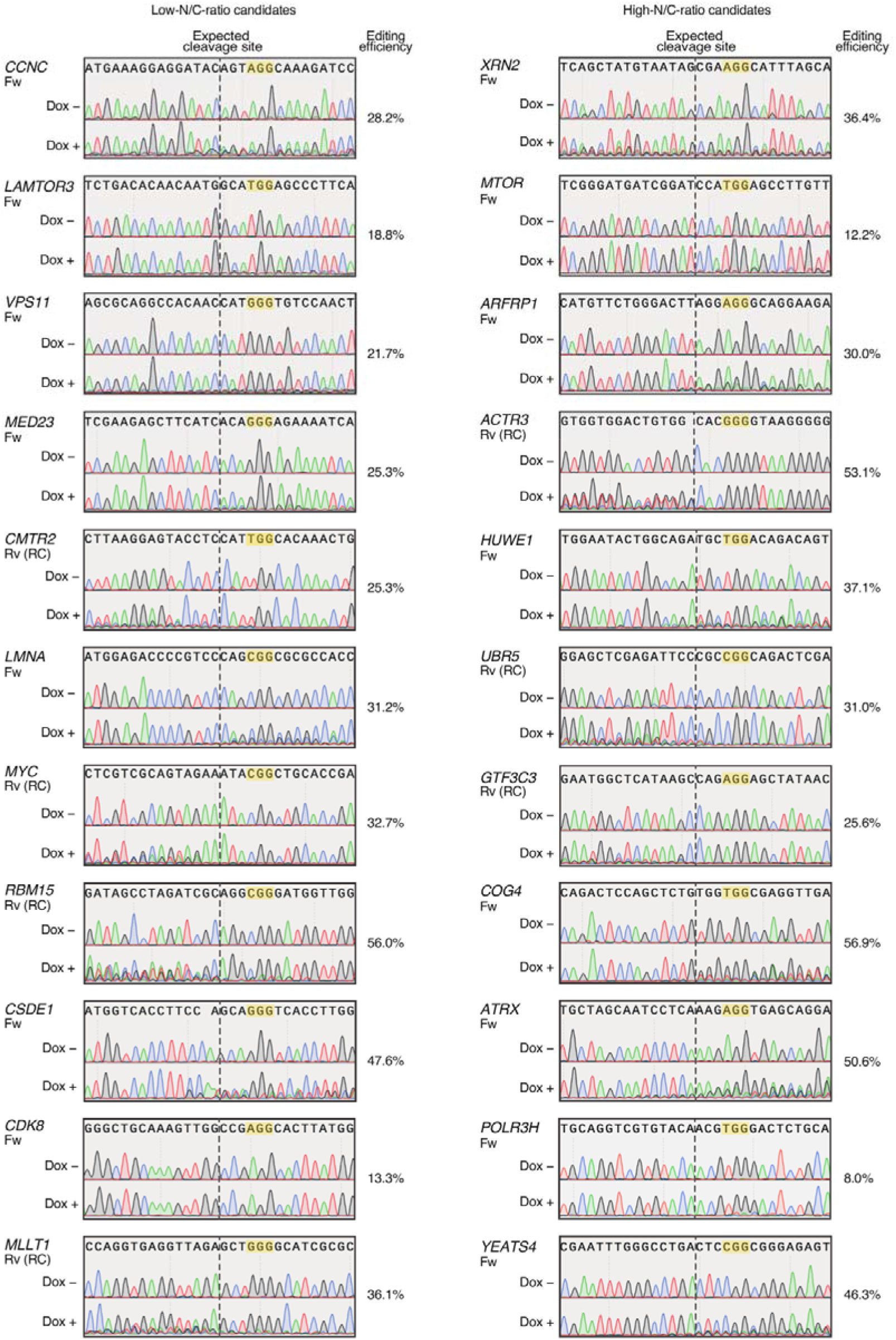
Sanger sequencing and TIDE analysis of the 22 candidate genes selected for validation. Low- and high-N/C-ratio candidates identified in the genome-wide CRISPR screen are shown in the left and right columns, respectively. For each gene, Tet::Cas9 HeLa cells were cultured in the absence (Dox−) or presence (Dox+) of doxycycline for 7 days, and the CRISPR-targeted regions were analysed by Sanger sequencing. PAM sequences are highlighted in yellow, and vertical dashed lines indicate the expected SpCas9 cleavage sites. Editing efficiency was estimated from Sanger sequencing chromatograms using TIDE. The sequencing primers used for each gene is indicated as Fw (forward) or Rv (reverse). Reverse-primer reads that were reverse-complemented for display are indicated as Rv (RC).

**Fig. S4.**
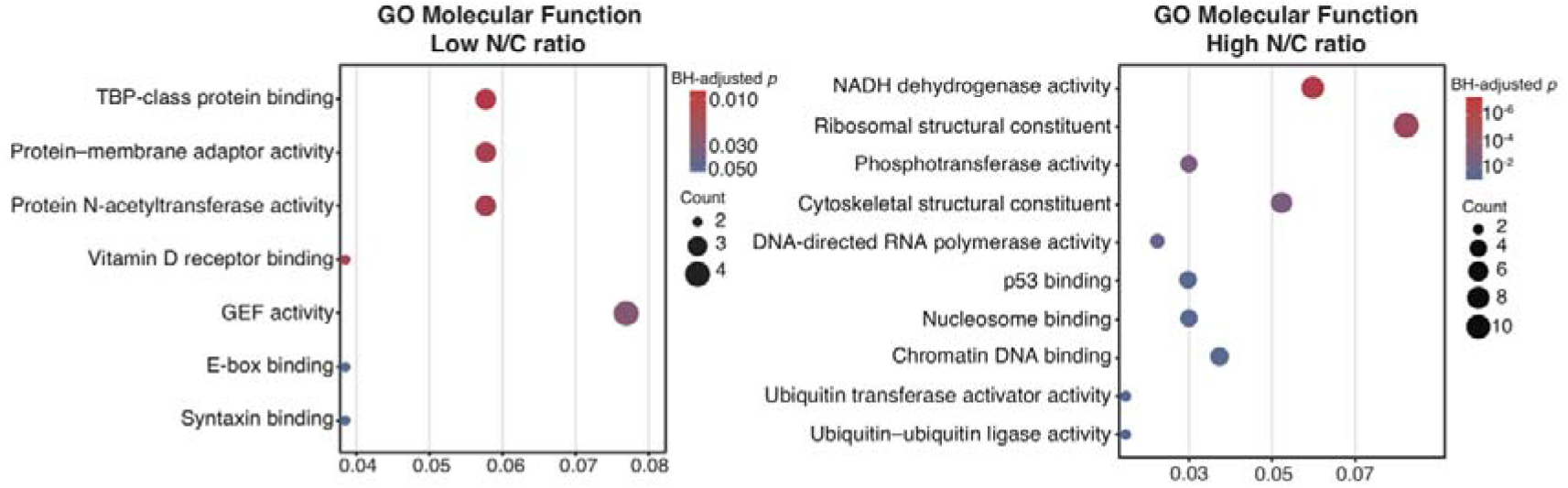
GO molecular function enrichment analysis of genes enriched in low- and high-N/C ratio populations. GO Molecular Function (MF) enrichment analysis of genes enriched according to the CRISPR screen in low- (left) and high-N/C ratio (right) populations. Dot size indicates the number of genes associated with each term, and colour indicates the Benjamini–Hochberg-adjusted *p*-value.

**Figure S5.**
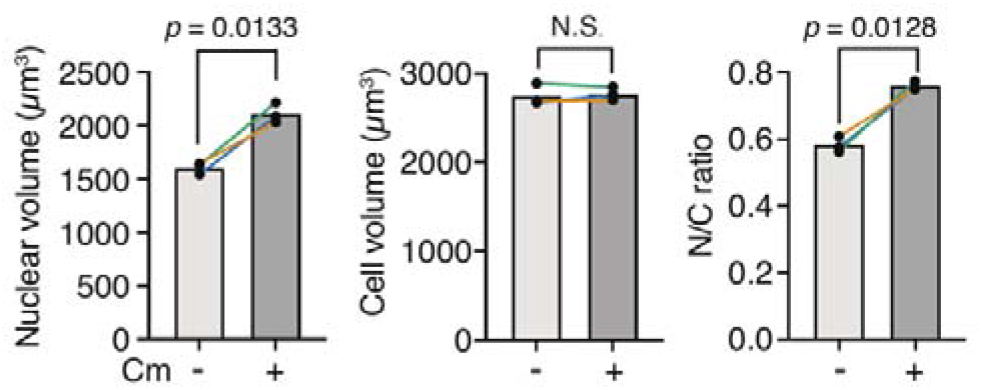
Chloramphenicol treatment increases nuclear volume and the N/C ratio in RPE-1 cells. In RPE-1 cells, nuclear volume (left), cell volume (middle), and N/C ratio (right) were measured 24 h after treatment with mitochondrial translation inhibitor chloramphenicol (Cm). At least 1,000 cells were quantified in each replicate. Each dot represents an independent experiment. Lines connect data points obtained from the same experiment. *p*-values were assessed using a two-tailed paired Student’s t-test. Error bars indicate SD. n = 3 biological replicates.

**Figure S6.**
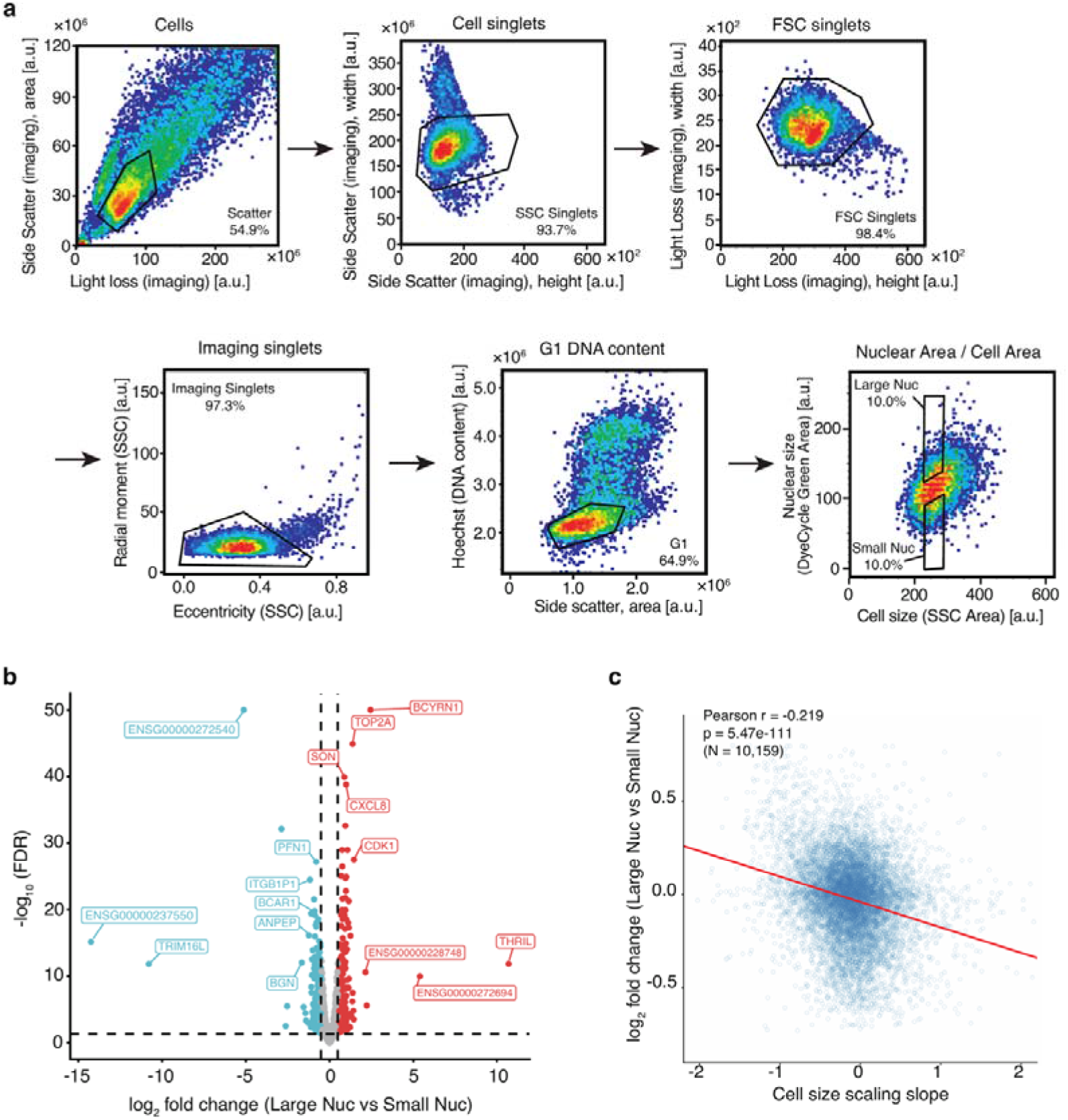
Transcriptomics analysis of RPE-1 cell sorted by the nuclear size. **a**, Sequential gating strategy used for image-enabled cell sorting. First, cells were selected on the basis of side scatter area and light loss area, followed by gating for SSC singlets and FSC singlets using imaging-derived pulse width and height parameters. Regular shaped singlets were further selected based on SSC eccentricity and radial moment. G1-phase cells were then identified by Hoechst-stained DNA content. Within the gated G1 population, nuclear size (DyeCycle Green Image Area) was plotted against cell size (SSC Image Area), a narrow cell size gate was set, within which the top and bottom 10% of cells by the nuclear area were collected as Large Nucleus and Small Nucleus cell populations, respectively. **b,** Volcano plot showing differentially expressed genes across the full annotated gene set, including protein-coding genes, non-coding RNAs, and uncharacterized loci, between small- and large-nucleus populations. Thresholds were set at FDR < 0.05 and |log_2_ fold change| ≥ 0.5. Blue dots indicate genes significantly upregulated in the small-nucleus population, whereas red dots indicate genes significantly upregulated in the large-nucleus population. **c,** Correlation between nuclear size–dependent transcription changes and cell-size-dependent mRNA scaling slopes in RPE-1 cells. Published gene-specific mRNA scaling slopes were obtained from You *et al.*^50^ and matched to our RNA-seq differential expression results. Each point represents a gene (N = 10,159). The x-axis indicates the mRNA scaling slope (log_2_), and the y axis indicates differential expression between large- and small-nucleus conditions (log_2_ fold change, Large/Small). Pearson correlation: r = −0.219, *p* = 5.47 × 10^−111^. The red line indicates the linear regression fit.

**Figure S7.**
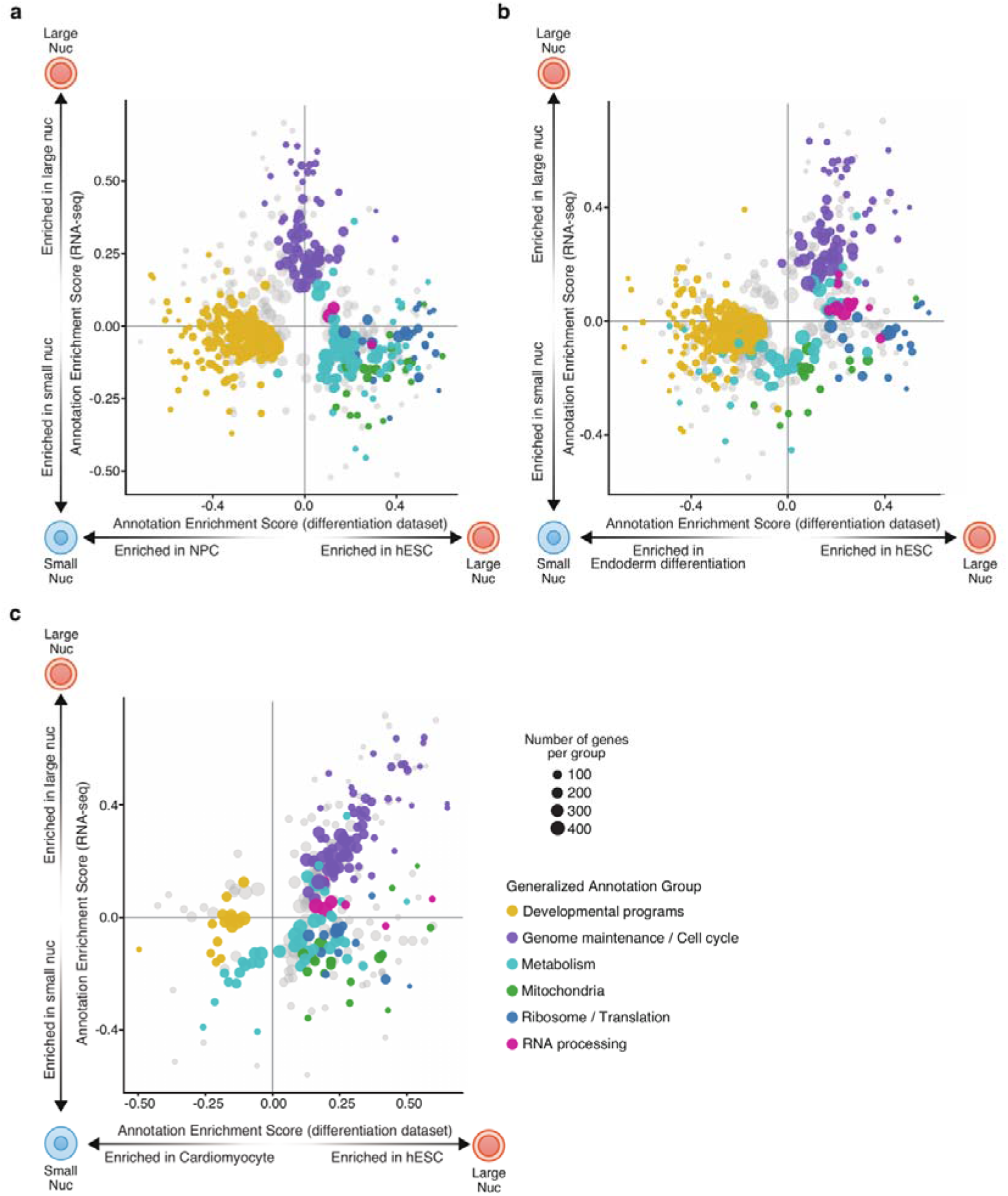
Comparative gene family enrichment analysis reveals similarity between the transcriptomes of small-nucleus RPE-1 cell and pluripotent stem cells undergoing differentiation. **a–c** 2D annotation enrichment analysis comparing transcriptional changes between small- and large-nucleus populations with gene expression changes in differentiation datasets (a, NPC versus hESC, GSE75748; b, definitive endoderm versus hESC, GSE109658; c, cardiomyocyte versus hESC, GSE85331). Gene annotation groups with BH-adjusted FDR < 0.02 are shown. Each dot represents an annotated gene group, with size indicating the number of genes. The x-axis indicates enrichment in the differentiation dataset, and the y-axis indicates enrichment in RNA-seq data comparing large- and small-nucleus populations.

**Figure S8.**
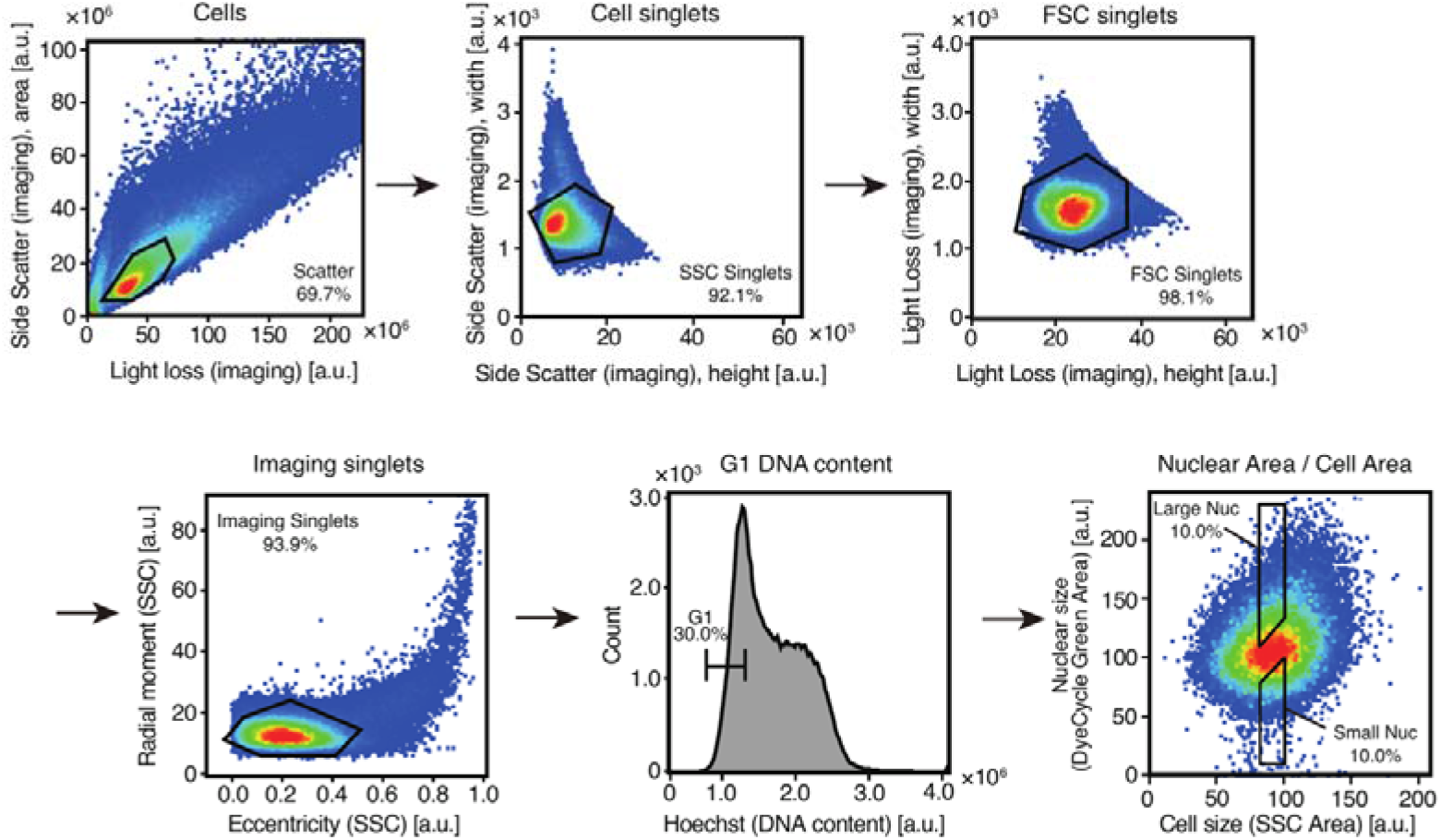
Sequential gating strategy for image-enabled sorting of mESCs according to their nuclear size. Cell singlets were first selected based on side-scatter area and light-loss area, followed by sequential gating of SSC singlets and FSC singlets using imaging-derived width and height parameters. Regularly shaped singlets were then selected based on SSC eccentricity and radial moment. G1-phase cells were identified based on Hoechst-stained DNA content. Within the gated G1 population, nuclear size, measured as DyeCycle Green image area, was plotted against cell size, measured as SSC image area. A narrow cell-size gate was applied, and the top and bottom 10% of cells according to nuclear area were collected as the Large Nucleus and Small Nucleus populations, respectively..

## Methods

### Cell culture and treatment

Non-transformed hTERT-immortalized human retinal pigment epithelium cells (cell line RPE-1) were obtained from the Stearns laboratory at Stanford. HeLa Tet::Cas9 cells (cTT20 cell line) were obtained from the laboratory of Iain Cheeseman^34^. RPE-1, HeLa, and HEK293T cells were maintained in Dulbecco’s modified Eagle medium with L-glutamine, 4.5 g/L glucose and sodium pyruvate (ThermoFisher # 41966052), supplemented with 10% heat-inactivated foetal bovine serum (FBS) and 1% penicillin/streptomycin. All cells were cultured at 37°C with 5% CO_2_. For drug treatments, cells were treated with chloramphenicol (50 μg/mL) and cycloheximide (10 μg/mL) for 24 h. Male naïve v6.5 mESCs (RRID: CVCL_C865), generously provided by Ali Shariati (University of California, Santa Cruz), were maintained on gelatin-coated plates in serum-based medium. Plates were coated with 0.1% (wt/vol) gelatin (G1890, Sigma-Aldrich) in Dulbecco’s phosphate-buffered saline (DPBS) for 30 min at 37 °C. The serum-based medium consisted of DMEM GlutaMAX (61965-026, Gibco) supplemented with 15% (vol/vol) ES-qualified FBS, 1% (vol/vol) MEM non-essential amino acids, 1% (vol/vol) sodium pyruvate, 1% (vol/vol) penicillin/streptomycin, 100 µM 2-mercaptoethanol and 10 ng/mL mouse leukemia inhibitory factor (LIF; Hyvönen Group, University of Cambridge). The medium was sterile-filtered using a Stericup Quick Release filtration system (S2GPU05RE, Millipore) before use.

### CRISPR/Cas9-mediated gene knockout

CRISPR/Cas9-mediated gene knockout was performed in Tet::Cas9 HeLa cells. For genome-wide screening, Tet::Cas9 HeLa cells were transduced with pooled lentiviral CRISPR/Cas9 sgRNA libraries targeting 18,408 protein-coding genes using five independent genome-wide libraries as previously described^33^. For validation experiments, individual sgRNAs targeting selected candidate genes were cloned into the CROP-seq-opti vector (Addgene #106280; BsmBI sites)^84^ and packaged into lentivirus in HEK293T cells using VSV-G and dr8.74 plasmids and TurboFect Transfection Reagent (Thermo Fisher Scientific #R0531). sgRNA sequences are listed in **Supplementary Data 10**. Cells were transduced with lentiviral supernatant in the presence of polybrene (10 μg/mL), selected with puromycin (2 μg/mL), and expanded for three days prior to Dox induction. Cas9 expression was induced by doxycycline (2 μg/mL) for 7 days prior to downstream assays. During culturing, HeLa cells were never allowed to grow above 60% confluency to prevent clumping.

### High-speed fluorescence image–enabled cell sorting for CRISPR screening

Live cells at a confluence of about 50% were detached from dishes using trypsin, resuspended in PBS at cell density of 10 million cells per mL, and stained with Vybrant DyeCycle Green (Thermo Fisher Scientific, #V35004; final concentration 10 µM) for 30 min at 37 °C. After staining, the cells were centrifuged for 3 min at 300g and resuspended at 20 million cells per mL in PBS containing 50 µg/ml DAPI as a cell viability dye. High-speed fluorescence image–enabled cell sorting was performed on a BD FACSDiscover™ S8 Cell Sorter (BD Biosciences)^33^. The gating strategy and image-based sorting parameters are shown in **Fig. S1a**. G1 cells were selected based on DNA content measured by DyeCycle Green fluorescence intensity. Nuclear size was quantified from acquired fluorescence images, and cell size was measured using SSC images. Based on these parameters, cells were sorted into high-, medium-, and low-N/C ratio populations. 10-15% of cells from the parental population were sorted into each of these three bins, resulting in about 2.2 million cells per bin.

### Genomic DNA isolation, library preparation and sequencing

Genomic DNA purification, sgRNA amplicon library preparation, and sequencing were performed following the CRISPR-screen amplicon-sequencing workflow described in Schraivogel et al., 2022^33^. Genomic DNA was prepared from sorted low, medium, and high N/C ratio bins, parental gate (G1) and unsorted HeLa samples using NEB Monarch genomic DNA purification kit (New England Biolabs), including the RNase treatment and elution in 100 µl elution buffer. gRNA loci were amplified from gDNA using a nested PCR strategy with outer primers (PCR1) and barcoded inner primers containing Illumina adapters (PCR2). Primer sequences and sample-to-i7 index assignments are provided in **Supplementary Data 11**.

PCR1 was done with up to 2 µg of gDNA, 1.5 µl 10 µM pU6 fwd, 1.5 µl 10 µM pLTR-CROP-rev primes, and 25 µl KAPA HiFi Hotstart Readymix in 50 µl total volume. For each of the three sorted bins from each of the five genome-wide sublibraries, six 50 µl reactions were set up to use up the total available amount of gDNA. Cycling conditions for PCR1 were one cycle at 95 °C for 3 min; 24 cycles at [98 °C for 20 s, 67 °C for 15 s, 72 °C for 15 s]; one cycle at 72 °C for 1 min; and cooling to 4 °C. PCR reactions of the same template were pooled and the product was purified with 0.8x AMPure XP (Beckman) with two 80% ethanol washes, and elution in 50 µl H_2_O.

PCR2 was done with 10 ng PCR1 product, 5 µl 3 µM CROPseq_libQC_i5_s1:6 staggered primer, 5 µl 3 µM CROPseq_i7:n barcoded primer, and 25 µl KAPA HiFi Hotstart Readymix in 50 µl total volume. Only one i5 staggered primer was used per reaction (i.e. no mixtures of staggered primers). Cycling conditions for PCR2 were one cycle at 95 °C for 3 min; 8 temperature cycles of 98 °C for 20 s, 67 °C for 15 s, and 72 °C for 15 s; one cycle at 72 °C for 1 min; and cooling to 4 °C. Product was pooled (by i7 barcode), purified as described for PCR1 and eluted in 40 µl H_2_O. Libraries were quality checked on a Bioanalyzer (Agilent) to yield a single product at around 300 bp. sgRNA library sequencing was performed by Novogene on Illumina Nextseq550 platform.

### sgRNA sequencing data processing and analysis

Following sgRNA Illumina sequencing, raw reads were processed with the nf-core/crisprseq v2.1.1 pipeline in screening mode, using Nextflow v24.04.2.5914^85,86^ and Mageck v0.5.9.5^87^. Guide library sequences were taken from the “Genome-wide library” tab of Table S4 in Schraivogel et al., 2022^33^. Sequences for sublibraries 1, 2, 3, 4, and 6 were included in the analysis (sublibrary 5 was excluded at the N/C sorting step due to the low transduction efficiency). Control guides included 351 non-targeting controls and 254 targeting controls. Raw counts from *Mageck count* (default parameters) were normalized, giving guide counts per gene as described in Schraivogel et al., 2022^33^. Specifically, raw sgRNA counts were divided by the median count of the targeting control sgRNAs in the corresponding sample to normalize differences in sequencing depth. These normalized counts were then scaled by multiplication with the median count of targeting controls across all samples.

To determine guides and associated genes with enrichment of reads in high and low gates we used MAUDE^35^ with normalized counts. The low bin was positioned between 0.001 and 0.12, medium bin 0.44 to 0.56 and high bin 0.88 to 0.999. The parent gate was used as the unsorted bin and all targeted and non-targeted controls were used. Mu optimization limits were -2 and 2. We used a *significanceZ* cutoff of <-10 or >10 and an FDR cut-off of 0.01 to call enriched genes.

### Assessment of genome editing efficiency in single-gene knockouts

Genome editing efficiency was evaluated in single-gene knockout non-clonal populations of Tet::Cas9 HeLa cells by comparing cells with and without Dox-induced Cas9 expression. Genomic DNA was extracted using the GeneJET Genomic DNA Purification Kit (Thermo Scientific, K0721). Target loci were amplified by PCR using primers flanking the sgRNA target sites. Sanger sequencing was performed on the PCR products, and editing efficiency was quantified using the TIDE (Tracking of Indels by DEcomposition) web tool (https://tide.nki.nl/)^88^.

### High-speed fluorescence image–enabled cell sorting for RNA sequencing, immunoblotting, and mESC differentiation assays

Live RPE-1 cells and mESCs were detached from dishes using trypsin, resuspended in culture medium, and co-stained with Hoechst 33342 (1:1000 dilution; Thermo Fisher Scientific, #62249) and Vybrant DyeCycle Green (1:1000 dilution; Thermo Fisher Scientific, #V35004; final concentration 5 µM) for 30 min at 37 °C. Cells were subjected to high-speed fluorescence image–enabled cell sorting on a BD FACSDiscover™ S8 Cell Sorter (BD Biosciences). The gating strategies for RPE-1 cells and mESCs are shown in **Fig. S6a** and **S8**, respectively.

### mESC differentiation assay and flow cytometry

Small- and large-nucleus mESC populations isolated by ICS were seeded on gelatin-coated plates in serum-based medium lacking LIF and supplemented with 1 µM retinoic acid (RA; HY-14649, MedChemExpress). After 36 h, cells were harvested by trypsinization and fixed with 3% formaldehyde in PBS for 10 min at 37 °C. Cells were then permeabilized with 90% ice-cold methanol in PBS for 30 min on ice. After washing with PBS, cells were blocked with 3% BSA in PBS for 30 min at 37 °C. Cells were co-incubated with rabbit monoclonal anti-NANOG [D2A3] (Cell Signaling Technology, Cat# 8822S; 1:20) and goat anti-SOX1 (Bio-Techne, Cat# AF3369; 1:20) in 3% BSA in PBS for 1 h at 37 °C. After washing with PBS containing 1% BSA and 0.05% Tween-20, cells were incubated with Alexa Fluor 568-conjugated donkey anti-rabbit IgG (H+L) (Invitrogen, Cat# A10042; 1:1,000) and Alexa Fluor 488-conjugated donkey anti-goat IgG (H+L) (Invitrogen, Cat# A11055; 1:1,000) in 3% BSA in PBS for 30 min at 37 °C. Cells were washed and analyzed using an Attune NxT flow cytometer (A24858, Invitrogen).

### RNA extraction and sequencing

Total RNA was extracted using the Direct-zol RNA Microprep kit (Zymo Research, #R2061) according to the manufacturer’s instructions. RNA quality and concentration were assessed prior to library preparation. Poly(A)-selected, non-directional libraries were prepared and sequenced (PE150) by Novogene using the Illumina NovaSeq X Plus platform.

### RNA-seq data processing

Raw RNA-seq reads were adapter-trimmed and quality-filtered using fastp (v1.1.0). Transcript abundances were quantified with Salmon (v1.10.3) in quasi-mapping mode against the GENCODE v45 human transcriptome (GRCh38), with sequence-specific and GC bias correction enabled. Transcript-level estimates were summarized to gene-level counts using tximport (v1.36.1). Differential expression analysis was performed in R (v4.5.0) using DESeq2 (v1.48.2). Gene expression was modeled using the design formula ∼ day + condition to control for replicate-specific variability while estimating the effect of nuclear size. Log_2_ fold changes were shrunk using apeglm (v1.30.0) to improve effect size estimation. Genes with an adjusted p-value (false discovery rate, FDR) < 0.05 and an absolute log_2_ fold change ≥ 0.5 were considered differentially expressed. Genes with NA adjusted p-values were excluded due to independent filtering of low-count features.

### Estimation of nuclear volume by fluorescence microscopy

Cells plated on glass-bottom 2-well chambered slides (µ-Slide, Ibidi) were stained with Hoechst 33342 (Thermo Fisher Scientific, #62249), which was added directly to the culture medium at a 1:6000 dilution, and incubated at 37 °C for 10 min. Fluorescence images were acquired using a ZEISS Axio Observer 7 epifluorescence microscope equipped with an Axiocam 820 mono camera and controlled by ZEN 3.10 (blue edition). Nuclear area was measured in Fiji (ImageJ) by automated segmentation of Hoechst fluorescence images using automated IsoData thresholding followed by watershed separation to distinguish touching nuclei. Objects touching the image border were excluded from analysis. Nuclear volume was estimated from the two-dimensional nuclear area (A) measured from microscopy images by assuming a spherical nuclear geometry. The nuclear radius (r) was calculated from the measured nuclear area according to the following relationship:

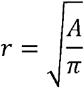

Based on this radius, nuclear volume (V) was calculated assuming a spherical shape:

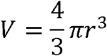

### Cell size measurement by Coulter counter

After nuclear size measurement, the cells were harvested from the chambered for a subsequent cell volume measurement. Cells were washed with PBS, detached with trypsin, and resuspended in culture medium. The entire cell suspension was diluted in 12 mL of ISOTON II diluent (Beckman Coulter) in an Accuvette. Cell size distributions were measured using a Multisizer 4e Coulter Counter (Beckman Coulter) and analysed according to the manufacturer’s instructions.

### Quantification of nuclear and cytoplasmic protein and RNA distribution

To quantify the subcellular distribution of total cellular protein and RNA, fluorescence imaging–based analysis was performed. Cells were cultured on glass-bottom dishes. For protein analysis, live cells were stained with the amine-reactive fluorescent dye carboxyfluorescein succinimidyl ester (CFSE) (Cytek Biosciences, 132-0850-U500, final concentration 5 µM), together with Hoechst 33342 (1:6,000 dilution; Thermo Fisher Scientific, #62249). For RNA analysis, cells were fixed with cold methanol for 10 min, washed with PBS, and stained with SYTO RNASelect Green Fluorescent Cell Stain (Invitrogen, S32703, final concentration 500 nM) for 20 min at room temperature, followed by Hoechst 33342 staining as described above. Fluorescence images were acquired, and nuclear regions were defined based on the Hoechst signal. Whole-cell regions were defined using the protein or RNA signal, and cytoplasmic regions were defined as the whole-cell area excluding the nuclear region. For each cell, total fluorescence intensity was quantified separately within the nuclear and cytoplasmic regions.

### Image-enabled immunofluorescent flow cytometry

Cells were harvested by trypsinization and fixed in 3% formaldehyde in PBS for 10 min at 37 °C. Samples were then permeabilized with 90% ice-cold methanol/PBS and incubated on ice for 30 min. After washing with PBS, cells were blocked in 3% BSA in PBS for 30 min at 37 °C and then incubated with the appropriate primary antibodies in 3% BSA in PBS for 1 h at 37 °C. The following primary antibodies were used: anti-trimethyl-Histone H3 (Lys27) [C36B11] (Cat# 9733, 1:200) and anti-Histone H3 [1B1B2] (Cat# 14269, 1:400) (Cell Signaling Technology). After washing with PBS containing 1% BSA and 0.05% Tween-20, cells were incubated with species-specific fluorophore-conjugated secondary antibodies (1:1000 in 3% BSA in PBS) for 30 min at 37 °C. The following secondary antibodies were used: Alexa Fluor 647-conjugated donkey anti-mouse IgG (H+L) (Cat# A31571) and Alexa Fluor 568-conjugated donkey anti-rabbit IgG (H+L) (Cat# A10042) (Invitrogen). Cells were then co-stained with 4’,6-Diamidino-2-Phenylindole (DAPI; Thermo Fisher Scientific, Cat# 3571, 300 nM) and Vybrant DyeCycle Green (Thermo Fisher Scientific, Cat# V35004, final concentration 5 µM) for 10 min at room temperature. Flow cytometry was performed using a BD FACSDiscover S8 Cell Sorter. Secondary-antibody-only controls were prepared in parallel and stained with the corresponding fluorophore-conjugated secondary antibodies, DAPI, and Vybrant DyeCycle Green, but without primary antibodies. These controls were used to estimate nonspecific fluorescence background. Because secondary-antibody background fluorescence increased approximately linearly with cell size, background subtraction for each cell was performed using a linear fit of the background signal against cell size. For each fluorescence channel, secondary-antibody-only control cells were used to fit a linear regression model between SSC imaging-derived cell size and fluorescence signal. For each immunofluorescently stained cell, the expected background fluorescence was estimated from the fitted linear model using its SSC imaging size, and the predicted background value was subtracted from the measured fluorescence signal. Background-corrected fluorescence values were used for quantification and analysis.

### Immunoblotting

For immunoblotting, resuspended in ice-cold hypotonic buffer containing 10 mM Tris-HCl (pH 7.4), 10 mM KCl, 1.5 mM MgCl□, 0.05% Triton X-100, and 1 mM DTT, supplemented with protease inhibitor cocktail (Sigma-Aldrich, P8340). After incubation on ice for 10 min, the samples were briefly vortexed and centrifuged at 1,500 × g for 5 min at 4°C. The nuclear pellets were washed once with the same hypotonic buffer and centrifuged again at 1,500 × g for 5 min at 4°C. Histones were extracted from the nuclear pellets with 0.2 N HCl by overnight rotation at 4°C. Following centrifugation at 15,000 × g for 10 min at 4□°C, the supernatants containing acid-extracted histones were collected and neutralized with Tris-HCl (pH 7.4). Protein samples were separated by SDS–PAGE and transferred to nitrocellulose membranes. Membranes were blocked with 3% BSA in TBST and incubated overnight at 4 °C with primary antibodies diluted in the same buffer. The following primary antibodies were used: anti-H3K27me3 (Cell Signaling Technology, Cat# 9733) and anti-Histone H3 (Cell Signaling Technology, Cat# 14269). All primary antibodies were used at a dilution of 1:1000. Fluorescently labeled secondary antibodies were IRDye 680LT goat anti-mouse IgG (LI-COR Biosciences, Cat# 926-68020) and IRDye 800CW goat anti-rabbit IgG (LI-COR Biosciences, Cat# 926-32211), both used at a dilution of 1:10,000. Membranes were imaged using an Odyssey CLx Infrared Imaging System (LI-COR Biosciences) and analysed using Image Studio software (LI-COR Biosciences).

### LEXY-based optogenetic manipulation of nuclear protein localization

HeLa cells stably expressing an NLS–mScarlet3–LEXY probe were seeded on μ-Slide 4 Well chambers (ibidi, Cat# 80426). Prior to imaging, nuclei were stained with SPY650-DNA (Spirochrome, Cat# SC501) diluted 1:1000 in 1× PBS for 1 h in the dark. Fluorescence images were acquired using a ZEISS Axio Observer 7 epifluorescence microscope equipped with an Axiocam 820 mono camera and controlled by ZEN 3.10 (blue edition). To induce nuclear export of the LEXY probe, cells were exposed to 475-nm blue light for 10 s every 1 min, and mScarlet3 fluorescence images were acquired at 1-min intervals for 90 min. Nuclear export was quantified by measuring the decrease in nuclear mScarlet3 fluorescence intensity over time. After 90 min of blue light illumination, 475-nm illumination was stopped, and cells were imaged every 1 min for an additional 120 min to monitor nuclear re-import of the probe. Nuclear import was quantified by measuring the recovery of nuclear mScarlet3 fluorescence intensity over time. Nuclear regions were defined based on SPY650-DNA fluorescence signals, and nuclear mScarlet3 intensity was quantified within these regions. Background fluorescence was subtracted from all measurements prior to analysis. Fluorescence intensities were normalized to the initial nuclear intensity for export analysis and to the maximal recovered nuclear intensity for import analysis. The half-time (t1/2) of nuclear export and import was calculated using GraphPad Prism (version 11.0.1). The half-time (t1/2) of nuclear export and import was defined as the time required for nuclear fluorescence intensity to reach 50% of the change between the initial value and the plateau.

### Principal component analysis

For RNA-seq principal component analysis (PCA), raw RNA-seq count data were variance-stabilized using the vst function in DESeq2 with blind=FALSE. PCA was performed using the plotPCA function, with samples grouped by condition and pair.

### Gene Set Enrichment Analysis

Gene set enrichment analysis (GSEA) was performed to identify coordinated transcriptional programmes associated with N/C ratio differences. Genes were ranked according to shrunken log_2_ fold changes obtained from DESeq2 using apeglm shrinkage. Enrichment analysis was conducted using the fgsea R package (Bioconductor) against the MSigDB Hallmark gene sets (v2025.1.Hs), which represent curated and non-redundant biological states. Pathways with a false discovery rate (FDR) < 0.05 were considered statistically significant. GSEA was prioritized over over-representation approaches because it evaluates genome-wide expression patterns without relying on arbitrary differential expression thresholds.

### GO enrichment analysis of nuclear-size-dependent transcriptional changes

GO enrichment analysis of genes expressed at higher levels in small- and large-nucleus populations was performed using clusterProfiler in R with the human annotation database org.Hs.eg.db, with all genes detected in the RNA-seq dataset used as the background. GO:BP enrichment was tested using enrichGO with Benjamini–Hochberg multiple-testing correction. No significance threshold was applied to this analysis: terms were ranked by BH-adjusted *p*-value, and the seven highest-ranked terms for each population are displayed in **Fig. 5a**, with the ten highest-ranked terms for each population listed in **Supplementary Data 8** together with their adjusted *p*-values. This descriptive presentation was used because enrichment differed markedly in strength between the two populations: GO:BP terms enriched in large-nucleus cells reached high significance (BH-adjusted *p* < 10□²□), whereas no term enriched in small-nucleus cells reached conventional significance (lowest BH-adjusted *p* = 0.06), reflecting a transcriptional shift that is broadly distributed rather than concentrated in individual annotation terms.

### GO and CORUM enrichment analysis of CRISPR screen hits

GO enrichment analysis was performed separately for genes associated with low-N/C and high-N/C ratios using clusterProfiler in R with the human annotation database org.Hs.eg.db. All genes analysed in the CRISPR screen were used as the background. GO:BP and GO:MF enrichments were tested separately using enrichGO with Benjamini–Hochberg (BH) multiple-testing correction. Terms with P ≤ 0.1, BH-adjusted P ≤ 0.1, and q ≤ 0.1 were considered for downstream analysis. To reduce redundancy, enriched terms were ranked by statistical significance and assigned directly to a previously retained representative term when the overlap coefficient between their hit-gene sets was at least 0.70 for GO:BP and 0.50 for GO:MF. Representative terms were visualized using dot plots. Protein-complex enrichment analysis was performed separately for low-N/C and high-N/C hit genes using human complexes from the CORUM database, with all genes analysed in the CRISPR screen used as the background. Enrichment was tested using clusterProfiler, followed by BH multiple-testing correction. For visualization, complexes with a BH-adjusted P < 0.1 and at least two hit genes were retained. Redundant complexes were reduced using the same direct-to-representative approach based on overlap of their CORUM protein subunits, with an overlap coefficient cutoff of 0.50. Representative enriched complexes were visualized using dot plots.

### Integration of nuclear-size-dependent RNA-seq data with cell-size-dependent mRNA scaling slopes

Gene-specific cell-size-dependent mRNA scaling slopes in RPE-1 cells were obtained from You et al.^50^. These slopes describe how mRNA concentration changes as a function of cell size for each gene. Nuclear-size-dependent transcriptional changes in our RNA-seq dataset were quantified as DESeq2-derived log_2_ fold changes between large- and small-nucleus conditions, with positive values indicating higher expression in large-nucleus cells. Published mRNA scaling slopes were matched to our RNA-seq differential expression results by gene symbol. Analyses were restricted to genes with valid mRNA scaling slope values and valid log_2_ fold change values in both datasets. To reduce the influence of extreme outliers, genes were filtered to retain those within the 0.5th–99.5th percentile of nuclear-size-dependent log_2_ fold change values. Pearson correlation analysis was performed at the gene level between the published mRNA scaling slopes and the log_2_ fold changes from our RNA-seq dataset. The reported P value corresponds to the Pearson correlation test. Linear regression was used only to visualize the overall relationship between the two measurements.

### 2D annotation enrichment

Two-dimensional annotation enrichment analysis was performed to compare RNA-seq and CRISPR screen datasets, as described previously^89^. Genes detected in both datasets were used as the analysis universe and ranked by DESeq2 shrunken log_2_ fold change for RNA-seq and by Vesuvius significance Z score for the CRISPR screen. GO Biological Process gene sets were obtained from MSigDB using the msigdbr package, and human protein complexes were obtained from CORUM. GO:BP gene sets containing 15–500 genes and CORUM complexes containing 2–500 mapped subunits within the shared analysis universe were analysed. For each annotation group, enrichment scores were calculated from rank distributions along both axes by comparing the mean rank of genes or mapped subunits within the group with that of all remaining genes. Enrichment scores were tested using multivariate analysis of variance (MANOVA) on the rank values, followed by Benjamini–Hochberg correction applied within each annotation source. This analysis is used in two distinct ways. The comparison between the nuclear-size RNA-seq dataset and the differentiation datasets (**Fig. 5d** and **Fig. S7**) is significance-bearing: annotation groups with a source-specific FDR < 0.02 were considered significant and plotted, following the 2% false discovery rate recommended as the default by Cox and Mann, 2012^89^. The comparison between the CRISPR screen and the nuclear-size RNA-seq dataset (**Fig. 4g**) is exploratory and is used to display the relative positioning of functional categories across the two datasets rather than to establish significance. For that panel, a relaxed display threshold of source-specific FDR < 0.07 was applied, chosen so as to encompass the development- and differentiation-related categories discussed in the text; this is consistent with the BH-adjusted P ≤ 0.1 threshold used for exploratory term-level enrichment elsewhere in this study. The CORUM MYC–MAX-containing and Mediator-related complexes are additionally displayed irrespective of threshold, to facilitate comparison with **Fig. 2a**. No claim of statistical significance attaches to inclusion in **Fig. 4g**; enrichment scores and FDR values for all annotation groups tested are provided in **Supplementary Data 7**.

### Statistical analysis

Data represent means ± standard deviation (SD) or standard error of the mean (SEM), as indicated in the figure legends, from at least three independent experiments. The statistical significance of differences between mean values was tested using a two-tailed Student’s t-test or one-way ANOVA with Tukey’s multiple comparison test. For the N/C ratio, nuclear and cell volumes were measured from the same cell population using microscopy and a Coulter counter, respectively. The N/C ratio was defined as the ratio of the mean nuclear volume (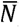) to the mean cell volume (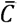). The standard error of mean (SEM) of the N/C ratio was estimated by error propagation from the SEM of nuclear volume (SEM*_N_*) and the SEM of cell volume (SEM_C_), assuming independence between nuclear and

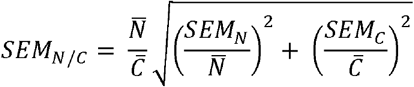

## Author Contributions

E.Z. and J.M.S. conceived the study. H.M., D.S., J.M.S. and E.Z. designed the experiments. E.Z. performed the CRISPR screen with assistance from D.S. A.J.R. performed bioinformatics analyses of the CRISPR screen data. R.S. performed and analysed mESC differentiation assays. H.M. performed and analysed all other experiments and curated the data. L.M.S., J.M.S. and E.Z. provided resources. H.M., J.M.S. and E.Z. wrote the manuscript with input from all co-authors. E.Z. supervised the study.

## Acknowledgements

We thank Richard Smith, Owen Gittins and Kenneth Jones for technical assistance; Shuyuan Zhang, Kurt Schmoller, Gabriel Neurohr, and all members of the Zatulovskiy and Skotheim laboratories for valuable discussions and feedback on this project; Iain Cheeseman for providing the cTT20 cell line and Ali Shariati for providing the mESCs. Image-enabled cell sorting for this project was performed on instruments in the EMBL Flow Cytometry Facility and University of Cambridge, Department of Pathology FACS Facility. We thank Beata Ramasz and the staff of both FACS facilities for their assistance. We thank Sandra Clauder-Münster for assistance with the EMBL-Stanford exchange visit, which led to the initiation of this collaborative project. This work was supported by the Medical Research Council (MR/X020290/1 Career Development Award to E.Z.), the NIH (P01 CA254867 grant to J.M.S.), and an Exchange Grant provided by the Life Science Alliance to E.Z.

